# Identification of a unique TUBB3^+^ cell population in the tuberculosis granuloma

**DOI:** 10.1101/2025.03.04.641465

**Authors:** Sarah C. Monard, Arnaud Métais, Maxime Pingret, Caroline Buscail, Maxime Caouaille, Marion Faucher, Serge Mazères, Renaud Poincloux, Yoann Rombouts, Bruce S. Klein, Marcel Wüthrich, Francisco J. Salguero, Simon Clark, Xavier Montagutelli, Etienne Simon-Lorière, Jean-Michel Molina, Nathalie de Castro, Nicolas Gaudenzio, Sergo Vashakidze, Claire Magnon, Cristina Vilaplana, Geanncarlo Lugo-Villarino, Christel Vérollet, Olivier Neyrolles

## Abstract

Tuberculosis (TB), caused by *Mycobacterium tuberculosis*, remains a leading cause of mortality worldwide. Granulomas, hallmark structures of TB in the lungs and other infected tissues, are critical sites of host-pathogen interactions, yet their full cellular composition is not completely understood. Here, we identify a previously unrecognized β3-tubulin (TUBB3)-positive cell population within TB granulomas in mice, guinea pigs, non-human primates, and TB patients. TUBB3 is a well-established pan-neuronal marker, yet these TUBB3^+^ cells are distinct from typical pulmonary resident cells and leukocytes. They exhibit a branched, elongated morphology, which is suggestive of neuron-like features. Intriguingly, their appearance is independent of adaptive immunity and is also observed in viral and fungal infections, but not in asthma. Our findings suggest the existence of a neuro-immune component within granulomas that may influence TB pathogenesis. Further investigation into the origin, function, and signaling pathways of these TUBB3+ cells is required to clarify their identity and potential role in host defense, which could reveal novel therapeutic targets for TB and other pulmonary infections.

## Introduction

Tuberculosis (TB), caused by the respiratory bacterial pathogen *Mycobacterium tuberculosis*, remains the leading cause of death from a single infectious agent, surpassing HIV/AIDS and malaria (Global Tuberculosis Report 2024). Despite its declining prevalence in many developed countries, TB continues to pose a significant public health burden, particularly in low- and middle-income countries, where it claims approximately 1.5 million lives annually. The disease primarily affects the lungs, where immune responses culminate in the formation of granulomas—organized multicellular structures that not only contain the infection but also provide a niche for *M. tuberculosis* persistence. Granulomas are composed of a diverse array of immune cells, as well as non-immune stromal and structural cells (Flynn and Chan 2022). Yet, their precise cellular composition, dynamic interactions, and role in disease progression remain incompletely understood.

Understanding the interplay between host cells and *M. tuberculosis* within granulomas has been central to TB research, with much focus placed on immune mechanisms (Flynn and Chan 2022). However, the role of the nervous system, which is increasingly recognized as a critical regulator of immunity, remains largely unexplored in TB. This gap in knowledge is particularly surprising given that the lungs, as the primary site of TB infection, are among the most innervated organs in the body (Blake et al. 2019). Sensory and autonomic nerves mediate essential physiological functions in the lungs, including bronchoconstriction, mucus secretion, and protective reflexes like coughing (Audrit et al. 2017). In the context of infections, these nerves not only detect pathogens but also influence local immunity through bidirectional communication with immune cells (Deng and Chiu 2021; Naqvi et al. 2023; Azzoni et al. 2024). This neuro-immune crosstalk, while documented in conditions like cancer and fibrosis, has received limited attention in infectious diseases, including TB.

Topical studies have begun to shed light on the complex interactions between the peripheral nervous system (PNS) and the immune system that communicate through a network of electrical and biochemical signals. Neurons not only express cytokine receptors but also produce cytokines, while immune cells express receptors for neurotransmitters and neuropeptides and can secrete neurotransmitters, facilitating cross-talk between the two systems (Boahen et al. 2023). During inflammatory conditions, the PNS undergoes significant remodeling, including axonogenesis (*i.e.*, the formation of new axonal branches) and, less frequently, neurogenesis (*i.e.*, the generation of new neurons) (Ayala et al. 2008; Magnon et al. 2013; Mauffrey et al. 2019; Gao et al. 2021; Ferdoushi et al. 2021). These processes are thought to influence the tissue microenvironment and modulate immune responses, yet their implications in the context of infectious diseases are poorly understood.

In TB, the co-evolution of *M. tuberculosis* and its human host has led to complex interactions between the pathogen and host systems, including the PNS (Alaniz et al. 1999; Barrios-Payán et al. 2016; Islas-Weinstein et al. 2021). For instance, *M. tuberculosis*-induced coughing, a hallmark symptom of TB, is not merely a mechanical reflex but also involves neural pathways modulated by the infection (Ruhl et al. 2020). However, whether and how the PNS contributes to granuloma formation and immune regulation in TB remains an open question. Addressing this knowledge gap could provide valuable insights into TB pathophysiology and uncover novel therapeutic targets.

In this study, we provide evidence for the involvement of the PNS in TB by identifying a novel population of cells positive for β3-tubulin (TUBB3), a pan-neuronal marker, within the lungs of different *M. tuberculosis*-infected hosts, including mice, guinea pigs, non-human primates (NHPs) and TB patients. Immunohistological analysis revealed increased bronchial innervation in infected lungs, alongside the appearance of TUBB3^+^ cells within granulomas and infection-induced lesions. These cells, which exhibit branching and elongation, are clearly distinguishable from bronchial innervation and do not express markers of known pulmonary cell types, suggesting a neuronal origin. Their localization near innate and adaptive immune cells raises intriguing questions about their potential role in immune modulation, particularly given that their emergence occurs independently of adaptive immunity and is observed in viral and fungal infections but absent in non-infectious inflammatory conditions like asthma. By uncovering a previously unrecognized neuro-immune component in TB granulomas, our findings provide new insights into granuloma biology and suggest potential avenues for therapeutic intervention to treat TB and other pulmonary infections.

## Results

### Immunohistological staining for TUBB3 reveals peripheral nervous system remodeling in the lungs of M. tuberculosis-infected mice

TUBB3 is the only tubulin isoform constitutively expressed in all mature neurons (Latremoliere et al. 2018). In the healthy lung tissue of mice, TUBB3 mainly decorate axons surrounding the bronchi and blood vessels (**Figure 1A**, images 1), where these axons support vital physiological functions such as constriction, dilatation and coughing (Audrit et al. 2017). At day 42 post-infection with the H37Rv strain of *M. tuberculosis*, we observed a marked increase in both the intensity and number of TUBB3^+^ axons in these regions (**Figure 1A**, images 3, and **Figure 1B**). This observation was further supported by 3D reconstruction of TUBB3-stained lungs after tissue clearing (**Supplemental Figure 1A**, **Movie 1**). These results suggest that axonogenesis may be part of the lung’s response to *M. tuberculosis* infection.

**Figure 1.**
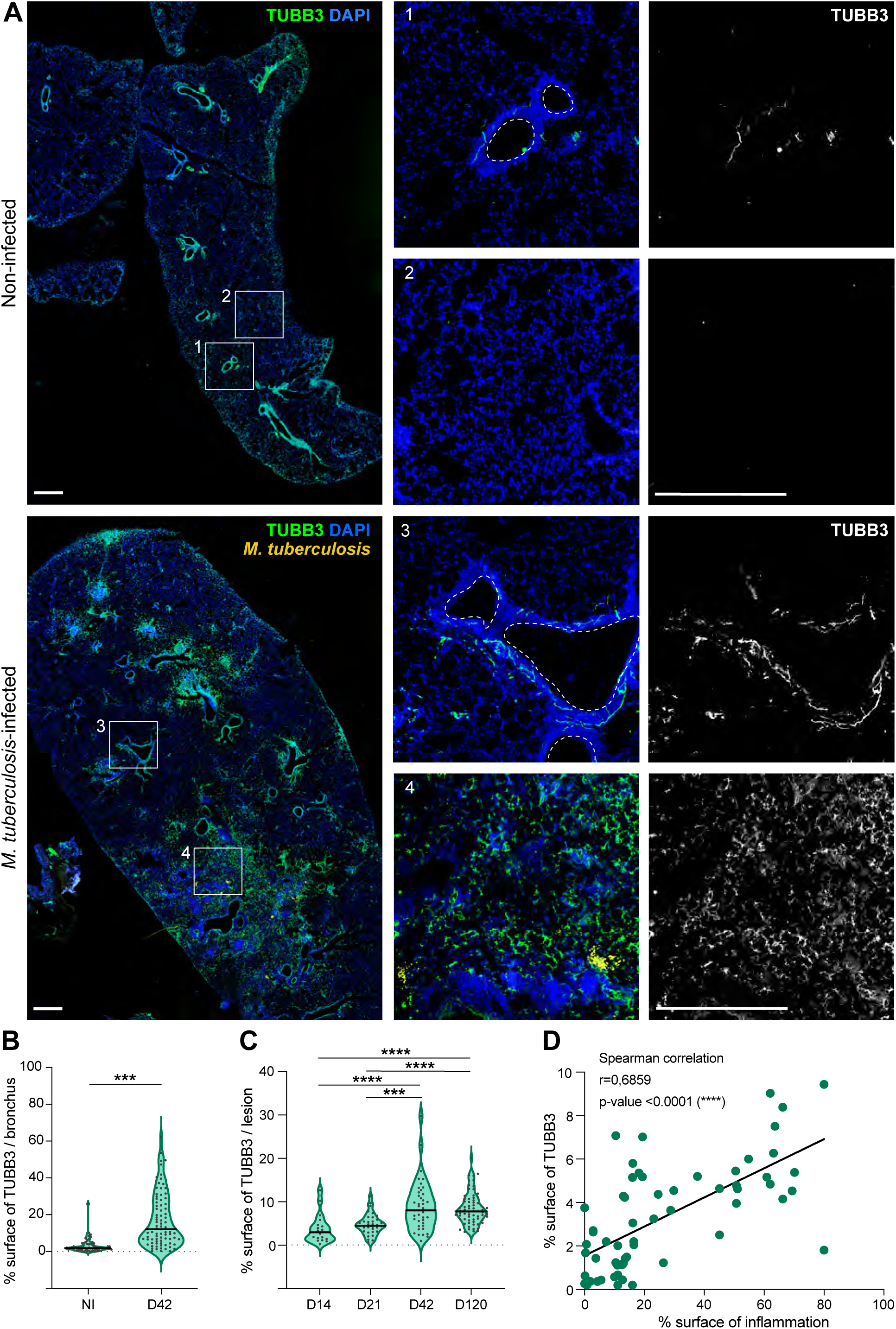
TUBB3^+^ cells accumulate in the lungs of *M. tuberculosis*-infected mice. **A.** Representative immunohistofluorescence images of lungs from non-infected (top) and *M. tuberculosis*-infected (bottom) mice at day 42 post-infection with the H37RV-dsRed strain (orange). TUBB3 (green) and nuclei (DAPI, blue) are shown. Images 1 and 3 are zoomed views of bronchi (dotted lines), and images 2 and 4 are zoomed views of the parenchyma. Data represent three independent experiments, with n=9 mice per condition. Scale bar, 500 µm. **B.** Percentage of TUBB3-stained surface per bronchia. Mann-Whitney test, three independent experiments, n=4 mice per timepoint. **C.** Percentage of TUBB3-stained surface per lesion in *M. tuberculosis*-infected mouse lungs. Kruskal-Wallis test with Dunn’s multiple group comparisons, three independent experiments, n=4 mice per timepoint. **D.** Correlation between the surface area of TUBB3 staining and the surface area of inflammation per lobe, defined by regions with dense number of nuclei. Spearman correlation. ***, p≤0.001; ****, p≤0.0001.

Surprisingly, we detected the appearance of numerous TUBB3^+^ cells within the parenchyma of *M. tuberculosis*-infected lungs (**Figure 1A**, image 4). These cells were completely absent in uninfected lungs (**Figure 1A**, image 2). TUBB3^+^ cells first became visible at day 21 post-infection, peaked at day 42 and plateaued until day 120, as shown by quantification of TUBB3^+^ surface areas per lobe and lesion (**Figure 1C**, **Supplemental Figure 1B-C**. The emergence of these cells coincided with infection-induced inflammation. Within the same lung lobes, TUBB3^+^ cells predominantly localized in inflamed areas, characterized by a high density of recruited cells, compared to non-inflamed regions (**Supplemental Figure 1D-E**). Notably, TUBB3^+^ cells were found near bacilli (**Supplemental Figure 1F**), but bacteria were never observed inside these cells, suggesting that TUBB3^+^ cells are not a direct target for this pathogen. Moreover, the extent of the TUBB3^+^ surface areas correlated significantly with the degree of inflammation, indicating that higher levels of inflammation are associated with a greater abundance of TUBB3^+^ cells (**Figure 1D**). Moreover, tissue clearing, and 3D reconstruction of the infected lung revealed that these parenchymal TUBB3^+^ cells are not connected to peribronchial TUBB3^+^ axons (**Supplemental Figure 1A** right, **Movie 1**).

Taken together, these findings demonstrate that *M. tuberculosis* infection triggers significant PNS remodeling in the lung, as evidenced by an increase in TUBB3^+^ axons surrounding the bronchi. Furthermore, we identified the emergence of previously unrecognized TUBB3^+^ cells within the lung parenchyma of infected mice.

### Appearance of TUBB3^+^ cells in the lung of M. tuberculosis-infected mice

Next, we focused on characterizing the atypical TUBB3^+^ cells induced by *M. tuberculosis.* High-resolution confocal microscopy confirmed that they contain nuclei (**Figure 2A**, **Movie 2**), further supporting that they are not extensions of preexisting axons, which typically have their cell bodies and nuclei within neuronal-specific ganglia outside the lung (Audrit et al. 2017). Parenchymal TUBB3^+^ cells are characterized by their branching structure and long cytoplasmic extensions, which are approximatively seven times the length of their nuclei (**Figure 2A-B**, **Movie 2**), facilitating connections with the surrounding cells. To determine whether TUBB3^+^ cells were capable of local proliferation, we analyzed Ki-67 expression by immunohistological staining in *M. tuberculosis*-infected murine lungs. Ki-67 is a protein present during all active phases of the cell cycle but absent in resting cells (Scholzen and Gerdes 2000). We did not detect Ki-67 expression in the nuclei of TUBB3^+^ cells, suggesting that they may be differentiated cells (**Figure 2C**, **Movie 3**). To evaluate the physiological and pathological relevance of this newly identified TUBB3^+^ cell population, we investigated its presence in Sp140-deficient mice, which are more sensitive to *M. tuberculosis* and develop key features of human pathology, such as the formation of hypoxic and necrotic granulomas (Ji et al. 2021). Indeed, the *Sp140* gene, located within the *Sst1* locus, represses type I interferon (IFN-I) transcription during bacterial infection. Consequently, *Sp140*^-/-^ mice more accurately replicate the hallmarks of human TB pathology. Infection with the *M. tuberculosis* Erdman strain of these mice also led to the abundant appearance of TUBB3^+^ cells in pulmonary lesions as observed at day 28 post-infection (**Figure 2D**).

**Figure 2.**
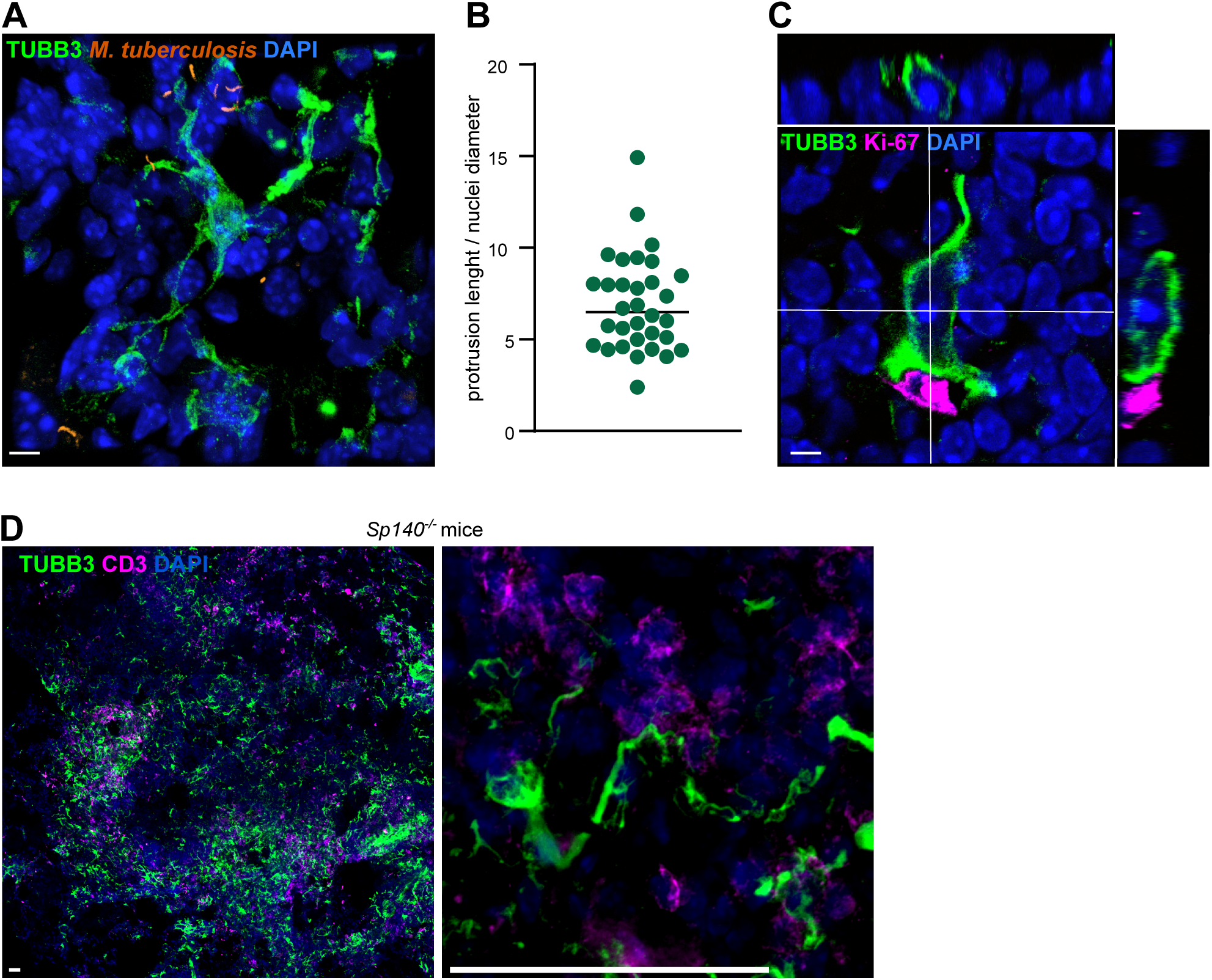
TUBB3^+^ cells exhibit a branched morphology and are not in a proliferative state. **A.** Representative immunohistofluorescence image of lungs from an *M. tuberculosis*-infected mouse at day 42 post-infection with the H37RV-dsRed strain (orange). TUBB3 (green) and nuclei (DAPI, blue) are shown. Data represent three independent experiments, with n=9 mice per condition. Scale bar, 5 µm. **B.** Quantification of the protrusion length relative to the nuclei diameter of TUBB^+^ cells. Thirteen TUBB3^+^ cells from seven different areas were analyzed, with each dot representing one protrusion. **C.** Representative immunohistofluorescence images showing that TUBB3^+^ cells (green) are not in a proliferative state, as indicated by the absence of Ki-67 staining (magenta), in the lungs of an *M. tuberculosis*-infected mouse at day 42 post-infection. Data represent n=2 mice. Scale bar, 5 µm. **D.** Representative immunohistofluorescence images of lungs of *M. tuberculosis*-infected *Sp140*^-/-^ mice at day 28 post-infection. TUBB3 (green), CD3 (T cells, magenta) and nuclei (DAPI, blue) are shown. Right panel shows the morphology of a TUBB3^+^ cell. Data represent two independent experiments, with n=3 mice. Scale bar, 50 μm.

While TUBB3 is widely recognized as a pan-neuronal marker, its expression in the lung has also been documented in non-neuronal cells. Indeed, TUBB3 is expressed in endothelial cells, neuroendocrine cells, and macrophages at steady state (Person et al. 2017; Mou et al. 2021), as well as in fibroblasts and cells undergoing epithelial-to-mesenchymal (EMT) or endothelial-to-mesenchymal (EndMT) transitions under pathological conditions (Duly et al. 2022). To determine the identity of the parenchymal TUBB3^+^ cells observed in *M. tuberculosis*-infected lungs, we conducted co-staining of TUBB3 with markers for epithelial/neuroendocrine cells (E-Cadherin), endothelial cells (CD31), fibroblasts (Vimentin), mesenchymal transition cells (α-smooth muscle actin, α-SMA), and immune cells (CD45) at day 42 post-infection (**Supplemental Figure 2A-E**). Our results showed that TUBB3^+^ cells do not colocalize with any of these markers, suggesting that they are not epithelial/neuroendocrine cells, endothelial cells, fibroblasts, cells undergoing EMT/EndMT, or immune cells.

As TUBB3^+^ cells do not express markers for typical pulmonary resident cell types and exhibit a branched and elongated morphology resembling neurons, we then investigated whether *M. tuberculosis*-induced TUBB3^+^ cells exhibit neuronal characteristics. While we detected both tyrosine hydroxylase (TH)^+^ autonomic sympathetic and vesicular acetylcholine transporter (VAChT)^+^ parasympathetic nerves around the bronchi of *M. tuberculosis*-infected murine lungs at day 42 post-infection (**Supplemental Figure 3A-B**, images 1), TUBB3^+^ cells did not colocalize with either of these markers (**Supplemental Figure 3A-B**, images 2). To explore the possibility that parenchymal TUBB3^+^ cells might be of sensory neuronal origin, we infected transient receptor potential vanilloid type 1 (TRPV1) reporter mice for 28 days. Sensory neuron axons were indeed present around the bronchi (**Supplemental Figure 3C**, images 1), however, we found no colocalization between TUBB3 and TRPV1 in the parenchyma (**Supplemental Figure 3C**, images 2), suggesting that these cells are not sensory neurons. This result was further supported by Na_V_1.8^+^ nociceptor depletion experiments using mice expressing diphtheria toxin fragment A (DTA) under the control of the Na_V_1.8 promoter (Abrahamsen et al. 2008). Specifically, we observed no significant changes in the appearance of parenchymal TUBB3^+^ cells in *M. tuberculosis*-induced lesions at day 28 post-infection in mice with or without Na_V_1.8^+^ nociceptors (**Supplemental Figure 4A**). Although weak, we were also able to detect a signal around the tracheae for Piezo2, specific for mechanoreceptors, however, no colocalization was observed with TUBB3^+^ cells in the parenchyma (data not shown).

While we could not conclude on the neuronal nature of the *M. tuberculosis*-induced TUBB3^+^ cells, we looked for the presence of progenitor cells expressing doublecortin (DCX) in *M. tuberculosis-*infected lungs using immunohistofluorescence analysis. Indeed, DCX^+^ brain-derived progenitor cells have been shown to migrate through the bloodstream, infiltrate tumors, and differentiate into TH^+^ nerve cells (Mauffrey et al. 2019). As shown in **Supplemental Figure 4B-D**, DCX^+^ cells were present in the inflamed areas surrounding blood vessels in *M. tuberculosis*-infected tissues, exhibited a classical small and round morphology of precursor cells, and co-expressed polysialylated neuronal cell adhesion molecule (PSA-NCAM), another marker for neuronal progenitors (Encinas et al. 2013), and the proliferation marker Ki-67. Given that DCX expression has also been reported in CD8^+^ T cells and microglia associated with amyloid-β plaques in Alzheimer’s disease (Unger et al. 2018), we performed co-staining of DCX and CD45 to exclude the possibility that these cells are leukocytes. Our results showed that approximately 80% of DCX^+^ cells do not express CD45, indicating that most of these cells are not leukocytes (**Supplemental Figure 4E**). These findings indicate that DCX^+^PSA-NCAM^+^ neuronal progenitor cells are present in the lungs during *M. tuberculosis* infection. However, the TUBB3^+^ cells observed in the infected lung do not express markers associated with sensory, sympathetic, or parasympathetic neurons.

Collectively, our results showed the emergence of TUBB3^+^ cells within the lung parenchyma of infected mice, which do not express markers of pulmonary cell populations, including sensory and autonomic neurons, or leukocytes. The emergence of parenchymal TUBB3^+^ cells is induced by the virulent *M. tuberculosis* strains H37Rv and Erdman in wild-type and *Sp140*^-/-^ mice, respectively. The observed peripheral nervous system remodeling in the lungs of *M. tuberculosis-*infected mice is also accompanied by the novel appearance of neuronal progenitor cells.

### The proximity of TUBB3^+^ cells to leukocytes suggests neuro-immune interactions during *M. tuberculosis* infection in mice

As TUBB3^+^ cells accumulate in inflamed lung areas during *M. tuberculosis* infection among infiltrating leukocytes, we analyzed their localization relative to specific immune cell types at day 42 post-infection. We found that these cells not only appear in *M. tuberculosis*-induced lesions, which are rich in F4/80^+^ macrophages and CD3^+^ T cells (**Supplemental Figure 5A**), but also in inducible bronchus-associated lymphoid tissue (iBALT), which contains B cells and follicular dendritic cells (**Supplemental Figure 5B**). iBALT, tertiary lymphoid organs that develop in the lung in response to infection, are typically found in the perivascular spaces near large blood vessels and airways (Silva-Sanchez and Randall 2020; Carpenter and Lu 2022). In both *M. tuberculosis*-induced lesions and iBALT, TUBB3^+^ cells were observed in proximity (∼3-5 µm of distance) to macrophages, CD4^+^ T cells, CD8^+^ T cells, and B cells (**Figure 3A-B**, **Supplemental Figure 5C-D**, **Movie 4**), suggesting that TUBB3^+^ cells are equally associated with these immune cell types. Next, to determine whether TUBB3^+^ cells might be involved in antigen presentation to T cells, we performed immunohistofluorescence for major histocompatibility complex class-I (MHC-I) and class-II (MHC-II). TUBB3^+^ cells expressed low levels of MHC-I and showed no detectable MHC-II expression (**Supplemental Figure 5E-F**), indicating that their primary function is unlikely to involve the presentation of peptide antigens.

**Figure 3.**
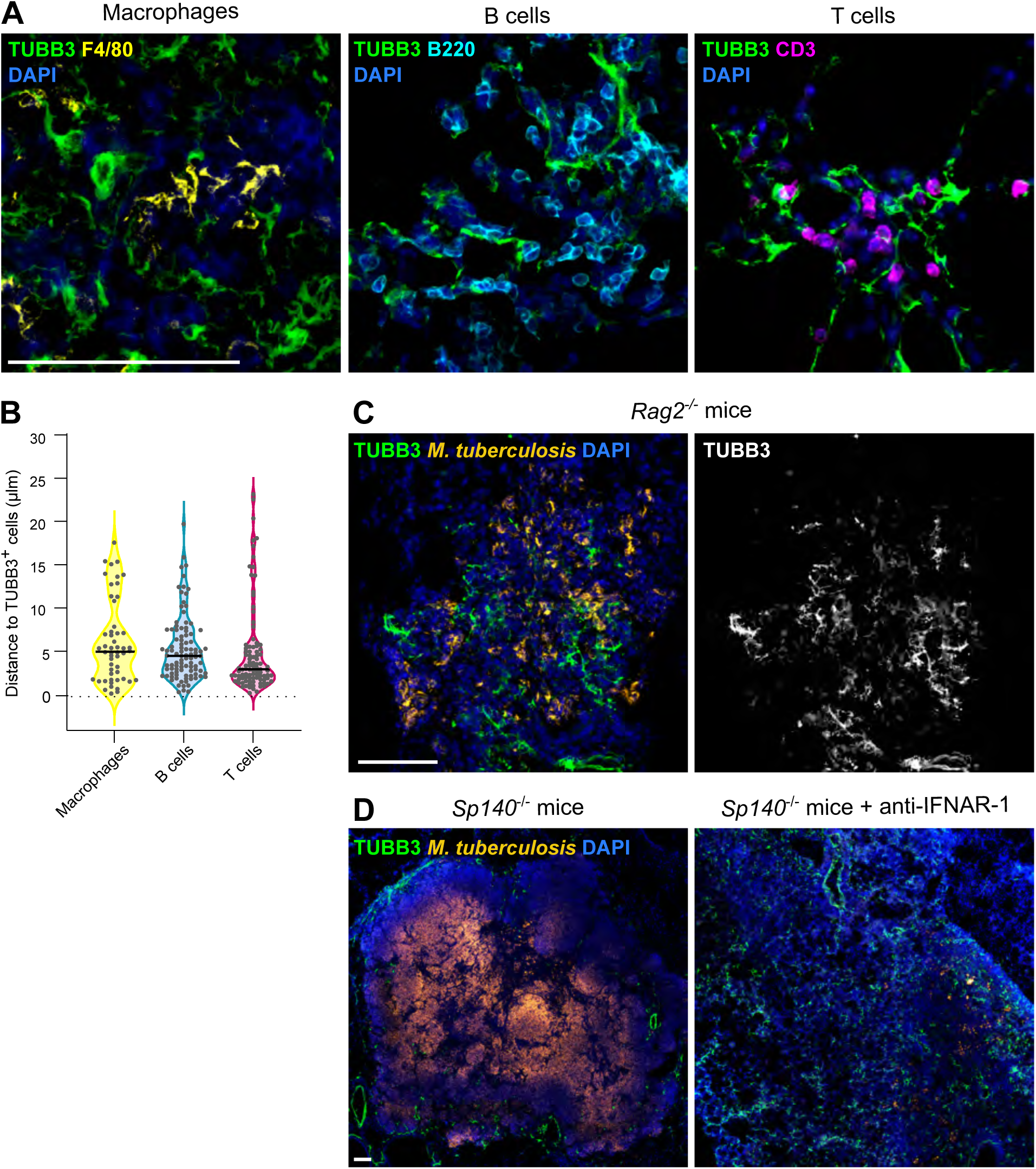
TUBB3^+^ cells are in close contact with immune cells during *M. tuberculosis* infection. **A.** Representative immunohistofluorescence images showing the proximity of TUBB3^+^ cells (green) to macrophages (F4/80, yellow, left), B cells (B220, cyan, middle) or T cells (CD3, magenta, right) in the lungs of *M. tuberculosis*-infected mice at day 42 post-infection. Nuclei (DAPI, blue) are shown. Data represent 2-3 independent experiments, with n=3 mice. Scale bar, 100 µm. **B.** Quantification of the distance between immune cells and TUBB3^+^ cells. Data were collected from at least three mice across 2-3 independent experiments. **C.** Representative immunohistofluorescence image showing TUBB3 staining (green) and nuclei (DAPI, blue) in *M. tuberculosis*-infected *Rag2*^-/-^ mice at day 28 post-infection with the H37Rv-dsRed strain (orange). Data represent two independent experiments, with n=6 mice. Scale bar, 100 µm. **D.** Representative immunohistofluorescence images of *M. tuberculosis* (Erdman-GFP strain, orange)-infected lungs of *Sp140*^-/-^ mice treated (right) or not (left) with anti-IFNAR-1 antibodies. TUBB3 (green) and nuclei (DAPI, blue) are shown. Data represent n=2 mice per group. Scale bar, 100 μm.

We then examined whether adaptive immunity and infection-associated cytokines, such as type-1 interferons (IFN-I) and IL17, contribute to the appearance of *M. tuberculosis*-induced TUBB3^+^ cells. To investigate the role of adaptive immune cells, we infected recombination activating gene 2 protein (Rag2)-deficient mice, which lack T and B cells. Notably, the absence of these immune cells did not prevent the appearance of TUBB3^+^ cells at day 28 post-infection (**Figure 3C**). Furthermore, as shown in **Figure 2D**, we found TUBB3^+^ cells in *Sp140*^-/-^ mice, which are susceptible to *M. tuberculosis* infection driven by elevated levels of IFN-I (Ji et al. 2021). In these mice, *M. tuberculosis* infection induces the formation of human-like granuloma structures (Ji et al. 2021), with TUBB3^+^ cells mainly localized at the granuloma periphery (**Figure 3D, left**). However, TUBB3^+^ cells were also abundant in infected mice treated with anti-IFNAR-1 (interferon-α/β receptor subunit 1) antibodies, a treatment that not only prevents TB lesion formation but also reduces pathogenicity, suggesting that IFN-I is not essential for their development (**Figure 3D, right)**. Finally, we evaluated the role of IL-17, which is involved in neuro-immune interactions such as in synaptic plasticity and pain induction (Ribeiro et al. 2019; Luo et al. 2021). Using *Il17a*-deficient mice, we found that this cytokine is not required for the presence of parenchymal TUBB3^+^ cells in *M. tuberculosis*-infected mice (**Supplemental Figure 6**). However, this does not exclude the potential involvement of other IL-17 isoforms in the emergence of TUBB3^+^ cells.

In summary, our results demonstrate that TUBB3^+^ cells are closely associated with immune cells in both *M. tuberculosis*-induced lesions and iBALT. Yet, they are unlikely to participate in antigen presentation to conventional T cell subsets. Moreover, their emergence in the lung during infection is independent of the adaptive immune system, as well as IFN-I and IL-17A signaling.

### TUBB3^+^ cells are distinctly localized within tuberculous granuloma in both animal models and humans

To evaluate the physiological and pathological relevance of this newly identified TUBB3^+^ cell population, we investigated its presence in TB granulomas across various animal models and in human patients. In the guinea pig model of infection, which replicates aspects of human TB pathology (Larenas-Muñoz et al. 2023a) and is commonly used to study coughing mechanisms (Bolser 2004), TUBB3^+^ cells were observed in the vicinity of TB granulomas, closely associated with macrophages and T cells (**Supplemental Figure 7**). *M. tuberculosis* infection in NHPs is considered the most accurate pre-clinical model for TB (Scanga and Flynn 2014). Like non-infected mice (**Figure 1**), under steady-state conditions, TUBB3 is located exclusively to axons surrounding the bronchi and blood vessels in the lung of NHPs (**Figure 4A**, left). In contrast, TUBB3^+^ cells were found in granuloma regions during both active and latent phases of TB in NHPs (**Figure 4A**, middle and right) (Kuroda et al. 2018). These *M. tuberculosis*-induced TUBB3^+^ cells are predominantly located in the outer layer of granulomas, near T cells and macrophages, reflecting the distribution observed in mice.

**Figure 4.**
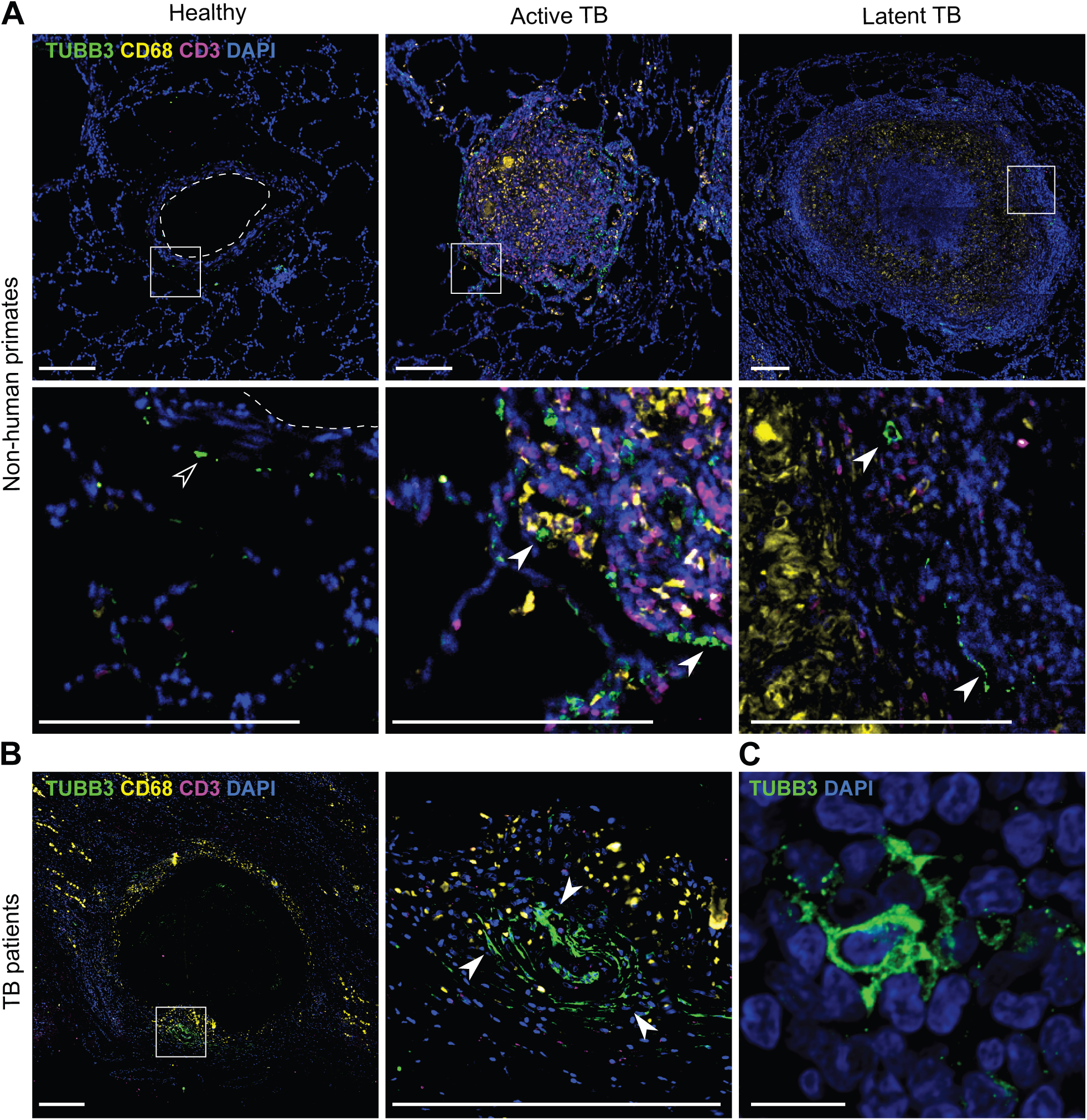
TUBB3^+^ cells are present in lung biopsies of *M. tuberculosis*-infected non-human primates (NHPs) and tuberculosis (TB) patients. **A.** Representative immunohistofluorescence images of lung biopsies from healthy NHPs (left) and NHPs infected with *M. tuberculosis* (active TB, left or latent TB, right). TUBB3 (green), CD68 (macrophages, yellow), CD3 (T cells, magenta) and nuclei (DAPI, blue) are shown. Empty arrowheads indicate peribronchial TUBB3^+^ axons; white arrowheads indicate parenchymal TUBB3^+^ cells; dotted lines outline the bronchi. n=3 individuals per group. Scale bar, 200 µm. **B.** Representative immunohistofluorescence images of a lung biopsy from a TB patient (Patient n°TB036, see **Table S1**). TUBB3 (green), CD68 (macrophages, yellow), CD3 (T cells, magenta) and nuclei (DAPI, blue) are shown. White arrowheads indicate parenchymal TUBB3^+^ cells. Scale bar, 200 µm. **C.** Representative immunohistofluorescence image of a TUBB3^+^ cell (green) containing a nucleus (DAPI, blue) in the lung granuloma from a TB patient (Patient n°TB011, see **Table S1**). Scale bar, 10 µm.

Even more importantly, we observed TUBB3^+^ cells in the vicinity of granulomas in lung biopsies from TB patients, where they were found near macrophages and T cells (**Figure 4B**, **Supplemental Figure 8A**). These TUBB3^+^ cells, which exhibit a morphology similar to those identified in mice (**Figure 4C** and **Figure 2A**), were absent in two diseased lung samples from non-TB patients but were present in between 2 and 24 areas within biopsies from all 11 TB patients analyzed (**Supplemental Figure 8A-B**). There were no statistically significant differences (P > 0.2) in the number of TUBB3^+^ areas between drug-sensitive TB (DS-TB) and multi-drug-resistant TB (MDR-TB) (**Supplemental Figure 8C**). Similarly, no significant differences were observed between patients with relapse and newly diagnosed TB cases (**Supplemental Figure 8D**) or between females and males (**Supplemental Figure 8E**). Of note, TUBB3^+^ cells were also detected in granulomas in lymph node biopsies from TB patients (**Supplemental Figure 8F**), albeit in much lower numbers than in the lungs. This suggests that these cells may represent a systemic host response to infection.

Taken together, these results show the appearance of these cells is conserved across species— including mice, guinea pigs, and NHPs—and occurs in both the lungs and lymph nodes of TB patients, highlighting their relevance in TB pathophysiology.

### Lung parenchymal TUBB3^+^ cells emerge during viral and fungal respiratory infections but are absent in asthma

Finally, we sought to determine whether the emergence of this novel lung parenchymal TUBB3^+^ cell population is specific to TB or represents a broader mechanism shared among pulmonary infections. To address this, we performed immunohistofluorescence to assess the presence of these cells in lung samples from mice with fungal and viral infections, as well as asthma. *Blastomyces dermatitis* and *Histoplasma capsulatum* are two fungal species responsible for granuloma formation in the lung (McBride et al. 2017). At 15 days post-infection, both fungi induced granulomas containing numerous TUBB3^+^ cells (**Figure 5A**). These cells displayed a branched morphology and were closely associated with CD3^+^ T cells (**Figure 5A**), resembling the pattern observed during *M. tuberculosis* infection. Similar TUBB3^+^ cells were identified in the lungs of mice infected with severe acute respiratory syndrome coronavirus 2 (SARS-CoV-2) on day 6 post-infection (**Figure 5B**), a time point characterized by high viral titers, significant weight loss, and moderate Th1/2/17 cytokine storm (Winkler et al. 2020; Oladunni et al. 2020). In this context, while TUBB3^+^ cells were located near T cells, they were diffusely distributed throughout the parenchyma rather than being clustered in inflamed areas. In contrast, parenchymal TUBB3^+^ cells were not detected in asthmatic lungs induced by intranasal challenge with house dust mites (HDM) extracts (**Figure 5C**) (Radermecker et al. 2019). Similar observations were made in a model of asthma based on intranasal challenge of *Alternaria alternata* extracts (data not shown).

**Figure 5.**
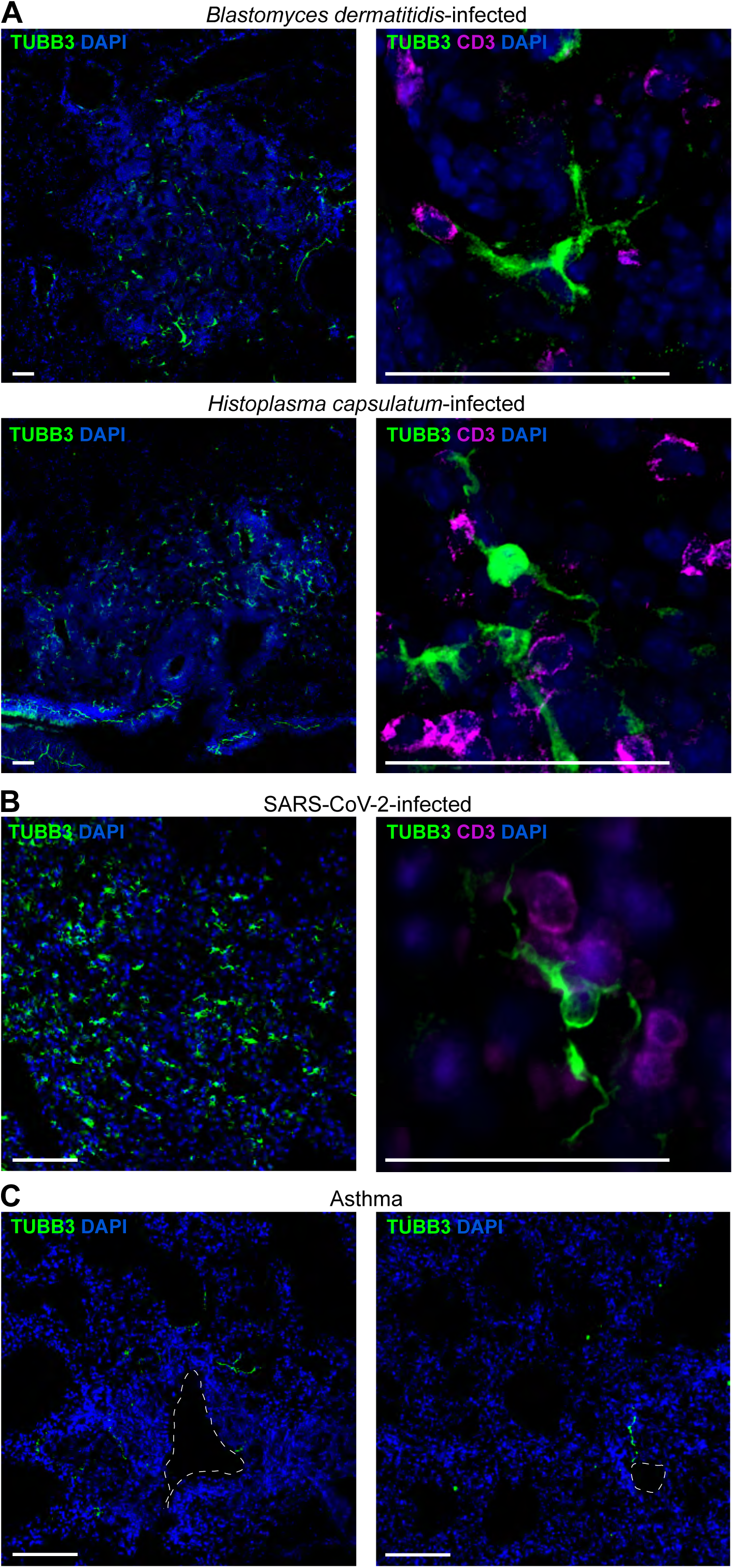
TUBB3^+^ cells accumulate in the lung during fungal and viral infections, but are absent in asthma. **A-B.** Representative immunohistofluorescence images of lungs from mice infected with *Blastomyces dermatitidis* (A, top) or *Histoplasma capsulatu*m (A, bottom) for 15 days, and with SARS-CoV-2 for 6 days (B). TUBB3 (green), CD3 (magenta, right) and nuclei (DAPI, blue) are shown. Data represent n=1-3 mice per timepoint. Scale bar, 200 µm. **C.** Representative immunohistofluorescence images of lungs from mice challenged with house dust mites (HDM) extracts and lipopolysaccharide as a model of asthma. TUBB3 (green) and nuclei (DAPI, blue) are shown. Dotted lines outline the bronchi. Data represent n=2 mice. Scale bar, 200 µm.

These findings suggest that the appearance of TUBB3^+^ cells is not specific to TB but may represent a broader host response to bacterial, viral and fungal infections, while being absent in non-pathogenic lung inflammation, such as asthma.

## Discussion

In this study, we explored potential PNS remodeling in the lung during *M. tuberculosis* infection. Our findings reveal a significant increase in TUBB3^+^ axons surrounding the bronchial tree in *M. tuberculosis*-infected murine lungs compared to non-infected controls. Additionally, we identify a novel TUBB3^+^ cell population within the lung parenchyma of *M. tuberculosis*-infected mice, guinea pigs, NHPs, and TB patients.

First, we report an increase in TUBB3^+^ axons in *M. tuberculosis*-infected murine lungs, suggesting that axonogenesis may occur in the lung during infection. Axonogenesis, or axonal sprouting, involves the growth of new axons from existing terminals and has been documented in both cancer and inflammatory disorders, such as lung fibrosis (Gao et al. 2021; Wu and Borniger 2023). Interestingly, the microenvironments of solid tumors and granulomas share several similarities, including secretion of both pro- and anti-inflammatory cytokines, hypoxia, central necrosis, and mechanisms of vascularization and tissue repair (Bickett and Karam, 2020; Ashenafi and Brighenti, 2022), suggesting that they may share mechanisms of axonogenesis induction (Magnon et al. 2013). One potential mechanism involves the secretion of extracellular vesicles, which are known to influence tumor innervation (Madeo et al. 2018) and are also secreted by *M. tuberculosis*-infected macrophages (Benet et al. 2021; Mehaffy et al. 2022). Interestingly, unlike in cancer, where axons infiltrate tumors, the axons induced by *M. tuberculosis* are mainly located around the bronchial tree without penetrating the tissue. As TUBB3 is a pan-neuronal marker, it is essential to determine whether this increase is specific to only sensory or autonomic neurons, or simply reflects a general rise in all lung innervation. Different neuronal subsets have distinct roles in sensory detection and signaling, bronchoconstriction, bronchodilation, coughing and mucus secretion (Audrit et al. 2017). Previous research by Ruhl *et al*. demonstrated that *M. tuberculosis* secretes sulpholipid-1, which activates nociceptors to induce coughing, thereby promoting bacillary dissemination (Ruhl et al. 2020). We infer that increased lung innervation may amplify the activation of the coughing reflex, further enhancing the secretion of infectious particles (Naqvi et al. 2023). Conversely, tissue innervation is also known to influence immune responses and disease progression (Schiller et al. 2021). In the context of TB, sympathetic neurons and catecholamines have been suggested to influence T helper cell differentiation (Barrios-Payán et al. 2016). Additionally, the cholinergic system increases the bacterial burden of *M. tuberculosis* while simultaneously suppressing pro-inflammatory cytokine production (Islas-Weinstein et al. 2021). Although these findings provide insight into the interplay between the autonomic nervous system and immunity to TB, further studies are needed to comprehensively evaluate the specific contributions of autonomic neurons in TB pathogenesis.

Second, we report the emergence of a novel TUBB3^+^ cell population in the lung parenchyma of *M. tuberculosis*-infected mice, guinea pigs, NHPs, and TB patients, underscoring the physiological relevance of this discovery. These cells are not connected to lung-innervating neurons and do not express typical markers of epithelial cells, endothelial cells, fibroblasts, muscle cells, or immune cells, highlighting the potential uniqueness of this cell population. TUBB3^+^ cells exhibit a branched morphology with long protrusions and a nucleus, which is suggestive of neuron-like features.

Neuron-like cells have been reported in non-neuronal tissues, particularly in the context of solid tumors, where they have been suggested to arise from the differentiation of migrating DCX^+^PSA-NCAM^+^ brain-derived progenitor cells (Mauffrey et al. 2019). Here, we observed the recruitment of DCX^+^PSA-NCAM^+^ cells to the inflamed areas surrounding blood vessels in *M. tuberculosis*-infected lungs. The localization of these cells suggests that they may be recruited from the bloodstream, although further studies are necessary to determine whether they originate from the brain. Although scarce, DCX^+^PSA-NCAM^+^ cells likely proliferate *in situ*, as indicated by their positivity for Ki-67, representing a potential precursor pool for the development of TUBB3^+^ cell during infection. For brain-derived progenitor cells to exit the brain, the blood-brain barrier (BBB) must be compromised. However, the impact of *M. tuberculosis* on BBB integrity remains a topic of debate, with conflicting findings reported in the literature (van Leeuwen et al. 2018; Proust et al. 2024), and further research being needed to clarify how progenitor cells might exit the brain in this setting.

To further investigate the potential neuron-like nature of *M. tuberculosis*-induced TUBB3^+^ cells, we co-stained TUBB3^+^ cells with markers specific to neuron subsets. We did not observe colocalization of TUBB3^+^ cells with markers for nociceptors, or for sympathetic and parasympathetic neurons. However, tissue fixation and staining for neuronal markers can be challenging, as previously noted in the brain, where specific tissue fixation methods, such as intra-aortic perfusion of paraformaldehyde before tissue harvesting, may be required to optimize the visualization of proteins expressed at low levels (Tu et al. 2021). Since we did not use this technique, we cannot definitively conclude whether TUBB3^+^ cells are related to neurons. Finally, to gain deeper insights into the identification of these cells, it is essential to go beyond immunohistofluorescence approaches that rely on single markers. Broader omics-based techniques should be employed to provide a comprehensive profile of this potential novel population. Such analyses would reveal whether *M. tuberculosis*-induced TUBB3^+^ cells are of neuronal origin, or if they possess unique, distinct features. This would also help determine whether these cells originate from the local differentiation of newly recruited neural progenitors or from the transdifferentiation of another cell type, as observed in other contexts (Mauffrey et al. 2019; Tang et al. 2022).

Third, we observed that parenchymal TUBB3^+^ cells are in close contact with immune cells, including macrophages, T cells and B cells, within *M. tuberculosis*-induced lesions and iBALT. Despite their proximity to immune cells, these TUBB3^+^ cells appear unlikely to directly present protein antigens to classical CD4 and CD8 T cells, as they express low levels of MHC-I and lack detectable MHC-II expression. However, it is possible that they may participate in the presentation of mycobacterial lipids via CD1-restricted molecules (Cohen et al. 2009), which warrants further investigation. At present, our data do not allow us to define the precise role of these cells in TB. Again, to get deeper insights into their function, omics-based analyses are essential for unraveling the key molecular pathways active in these cells. We hypothesize that physical interactions between immune cells and TUBB3^+^ cells may regulate immune cell activation, suppression or polarization, thereby shaping the lung microenvironment and its architecture. iBALTs, tertiary lymphoid structures that form in the lungs in response to chronic inflammation, infection, or other immune stimuli, are composed of various immune cells, including B cells, T cells, follicular dendritic cells, and macrophages (Dunlap et al. 2020; Carpenter and Lu 2022). These structures function similarly to organized lymphoid tissues by facilitating antigen presentation and lymphocyte activation, thereby contributing to local immune defense. In the context of TB, the formation of iBALT has been associated with the mycobacterial cell wall protein MmpL7 (Marin et al. 2019; Dunlap et al. 2020). Given the similarities in cell-to-cell interactions between TUBB3^+^ cells and immune cells within TB granulomas and iBALT, it is plausible that TUBB3^+^ cells play a regulatory role in the activation or repression of the immune response within iBALT. This interaction could potentially influence the development and function of these lymphoid structures.

Fourth, we describe the emergence of TUBB3^+^ cells in the vicinity of granulomas in *Sp140*-deficient mice, NHPs and TB patients, predominantly in the outer layer of the granuloma, a lymphocytes-enriched zone, much less pro-inflammatory than the inner granuloma core. It would be interesting to investigate whether TUBB3^+^ cells play a role in shaping granuloma architecture. Using a cohort with a limited number of TB patients, we observed no statistically significant differences in the occurrence of TUBB3^+^ cells in granulomas between DS-TB patients and MDR-TB patients, nor between patients with relapses and those with newly diagnosed TB. Despite these observations, expanding the cohort size may shed light into the role of TUBB3^+^ cells in TB physiopathology. Interestingly, we also observed the presence of TUBB3^+^ cells within lymph node granulomas of TB patients, while these cells were absent in the healthy regions of the same organs. This finding suggests that the emergence of TUBB3^+^ cells may not be confined to the lungs but could represent a broader systemic response to infection. Further investigation into their presence across different tissues could provide valuable insights into whether these TUBB3^+^ cells are consistently found in all *M. tuberculosis*-infected sites.

Last, we found that the emergence of TUBB3^+^ cells is not exclusive to *M. tuberculosis* infection. Similar cells were identified in close contact with T cells in the murine lung during infection with the virus SARS-CoV-2 and the fungi *B. dermatitidis* and *H. capsulatum*. SARS-CoV-2, which interacts with the nervous system, can cause neuroinflammation and a range of neurological symptoms, from temporary loss of taste and smell to persistent cognitive dysfunction, memory loss, and mood disorders characteristics of long COVID (Monje and Iwasaki 2022). The virus may enter the brain through various routes, including the olfactory pathway, vascular endothelium and BBB disruption, or transneuronal pathways from the lung (Rossi et al. 2022). If TUBB3^+^ cells are confirmed to be neuron-like cells, they may be involved in the brain-lung axis and potentially play a role in the pathology of long COVID. While parenchymal TUBB3^+^ cells were found in the lungs during bacterial, viral and fungal infections, they were notably absent in asthma models challenged with either HDM or *A. alternata* extracts. These results suggest that shared host response pathways to infection may drive the appearance of these cells. Immunity to infection primarily involves type-1 and type-3 immune responses, contrasting with allergic responses driven by type-2 immunity (Gopal et al. 2013; Lee and Lau 2017; Hammad and Lambrecht 2021; Segueni et al. 2021; Shen et al. 2023). We ruled out the necessity of adaptive immunity for the appearance of TUBB3^+^ cells, as these cells were still present in *Rag2*^-/-^ mice, which lack T and B cells. However, it is possible that innate immune cells or their associated molecular factors contribute to their emergence. Investigating the presence of TUBB3^+^ cells in mice deficient in T-box expressed in T cells (T-bet), GATA-binding protein 3 (GATA-3) and retinoic acid–related orphan receptor (ROR)γt, key transcription factors for type-1, -2 and -3 immunity, respectively, would help determine which immune response is most critical for their emergence.

In conclusion, our findings reveal that the PNS around the bronchial tree undergoes significant remodeling following *M. tuberculosis* infection, potentially altering the tissue microenvironment in ways that could either benefit the host or the pathogen. Additionally, we identified a novel population of branched cells expressing TUBB3, whose nature, origin, and function remain to be fully elucidated. This cell population consistently emerges in response to multiple strains of *M. tuberculosis*, predominantly in the vicinity of infection-induced granulomas in the lungs and lymph nodes and is conserved across host species. To our knowledge, TUBB3^+^ cells of this nature have not been documented in any organ during any infection. These cells may play a critical role in regulating the immune response to pulmonary infections. Their selective emergence, particularly in response to bacterial, viral, and fungal infections—but not in allergic inflammation such as asthma—suggests that they are either a key component of a broader host defense mechanism or a target of immune modulation by pathogens. Future research should focus on unraveling the molecular pathways driving the appearance of these cells and their potential contributions to immune regulation. Omics-based approaches will be instrumental in determining whether these cells represent a unique neuron-like population or a previously uncharacterized cell type with distinct features. Investigating their interactions with immune cells and their role in shaping the granuloma architecture could provide transformative insights into host-pathogen interactions. Ultimately, this work opens promising new avenues for understanding the complex interplay between the nervous and immune systems during infection.

## Material and Methods

### Human samples

#### Lung biopsies

The human lung tissue samples to be used for this study come from the collection C.0004270 obtained within the SH-TBL project in 2016; registered at the ClinicalTrials.gov database under code NCT02715271 and in the Biobank National Registry (Collections section) of the ISCIII and the Spanish Ministry of Science and Innovation. The SH-TBL project was reviewed and approved by both the Ethics Committee of the NCTLD (IRB00007705 NCTLD Georgia #1, IORG0006411) and the Germans Trias I Pujol Hospital Ethics Committee (EC: PI-16-171). Written informed consent was obtained for collection of biological material and data from all study participants before being enrolled. The samples were collected from TB patients undergoing therapeutic surgery. Details about the patients from whom lung biopsies were acquired for this study are annotated in **Table S1**. Symptom scores were assessed using the St. George’s Respiratory Questionnaire (SGRQ), a disease-specific tool designed to measure the impact of obstructive airway disease on overall health, daily life, and perceived well-being (Pasipanodya et al. 2007). SGRQ scores range from 0 to 100, with higher scores indicating greater limitations. All patients reported drinking alcohol less than once a month, and tested negative for human immunodeficiency virus, severe renal disease, immunosuppression and cirrhosis. Two lung samples from non-TB patients operated for bullosa pulmonary emphysema were included to the TB cohort in our study as potential controls for TB specificity. Human lung biopsies were provided as formalin-fixed paraffin-embedded (FFPE) samples under a material transfer agreement (MTA).

#### Lymph node biopsies

The human lymph node tissue samples used in this study were obtained from the Saint-Louis Hospital collection, under the supervision of Prs. Jean-Michel Molina, Philippe Manivet and Philippe Bertheau, as well as Dr. Nathalie de Castro. The samples were collected from TB patients undergoing therapeutic surgery, with written informed consent obtained from all participants prior to their enrollment for the use of frozen lymph node biopsies and associated data. Information on the patients from whom lymph node biopsies were acquired for this study are listed in **Table S2**. Human lymph node biopsies were both provided as FFPE samples under an MTA (BRIF number from the CRB LARIBOISIERE: B-0033-00064).

#### Non-human primate (NHP) samples

For this study, we used the NHP (Indian-origin Rhesus macaques, *Macaca mulatta*) model of acute tuberculosis. All animal procedures were approved by the Institutional Animal Care and Use Committee of Tulane University, New Orleans (LA), and were performed at the Tulane TNPRC under approval from IACUC (project numbers P3733, P3794, P3373 and P3628). They were performed in strict accordance with NIH guidelines. All NHPs were infected as previously described including the details regarding their physiopathology (Mehra et al. 2011; Foreman et al. 2016; Souriant et al. 2019). Briefly, aerosol infection was performed on NHPs using a low dose (25 CFU implanted) of *M. tuberculosis* CDC1551. Animals were euthanized when presenting four or more of the following conditions: (i) body temperatures consistently greater than 2 °F above pre infection values for three or more weeks in a row; (ii) 15% or more loss in body weight; (iii) serum CRP values higher than 10 mg/mL for three or more consecutive weeks, as it is a marker for systemic inflammation that exhibits a high degree of correlation with active TB in macaques (Mehra et al. 2011; Kaushal et al. 2012); (iv) CXR values higher than two on a scale of 0–4; (v) respiratory discomfort leading to vocalization; (vi) significant or complete loss of appetite; and (vii) detectable bacilli in BAL samples. NHP samples were provided as paraffin-embedded samples.

#### Mouse strains

All animal procedures adhered to French guidelines and were approved by the Ministry of Higher Education and Research (APAFIS agreement 34716 and 38001). C57BL/6 mice were purchased from Charles River. *Rag2^-/-^* mice were purchase from Jackson laboratory and bred at the Institute of Pharmacology and Structural Biology (IPBS) UMR 5089 (agreement F31555005). *Sp140*^-/-^ mice, kindly provided by Dr. Vance (UC Berkeley, United States), were also bred at IPBS. Control DTA, Nav1.8-DTA and TRPV1-YFP mice were provided by Dr. Gaudenzio and Dr. Lilian Basso (INFINITy, France), while *Il17a*^-/-^ mice were nicely provided by Dr. Gaya (CIML, France).

#### Bacteria

*M. tuberculosis* H37Rv, H37Rv-dsRed and Erdman strains were grown in suspension in Middlebrook 7H9 medium (BD) supplemented with 10% albumin-dextrose-catalase (ADC, BD), 0.05% Tween-80 (Sigma-Aldrich) and 100 µg/mL Hygromycin B (InvivoGen) (Lastrucci et al. 2015). For infection, growing *M. tuberculosis* at exponential phase was centrifuged at 3000 g and resuspended in Phosphate-Buffered Saline (PBS, Gibco) (MgCl_2_, CaCl_2_ free). Bacterial aggregates were dissociated by twenty passages through a 26G blunt needle. Bacterial suspension concentration was determined by measuring the optical density at 600 nm and then resuspended in PBS for *in vivo* infection.

#### Mice infections with H37Rv strain

Mice were anesthetized by intraperitoneal (i.*p*.) injections of ketamine and xylazine and were inoculated intranasally (*i.n.*) with 500-1,000 bacilli of H37Rv strain in 25 μL PBS. The inoculum dosage was confirmed by plating different dilutions on Middlebrook 7H11 OADC selective agar (Sigma-Aldrich). Plates were incubated at 37 °C and colonies were enumerated three weeks after.

#### SP140^-/-^ mice infection with Erdman strain and anti-IFNAR-1 treatment

*Sp140*^-/-^ mice were infected with 100 bacilli of Erdman strain using aerosol. The inoculum dosage was confirmed by plating lung extracts at day 1 post-infection on Middlebrook 7H11 OADC selective agar (Sigma-Aldrich). Plates were incubated at 37 °C and colonies were enumerated three weeks after. Anti-IFNAR-1 antibodies (Bioxcell, 500 µg) were injected via the *i.p.* route every other day from day 7 to 27 post-infection. Lung tissues of control or anti-IFNAR-1 treated mice were collected at day 28 post-infection.

#### Preparation of OCT-embedded tissues

Mice were euthanized at the appropriate time points after by an overdose of sodium pentobarbital via the *i.p.* route. Murine lung tissues were inflated by injection of 1 mL of PBS only or PBS with 2% agarose Ultrapure and collected (**Table S3**). Next, samples were fixed in 4% paraformaldehyde (PFA, Delta Microscopies), or in fixation buffer (0.05 M phosphate buffer, 0.1 M L-lysine (Sigma-Aldrich), 2 mg/mL NaIO_4_ (ThermoFisher Scientific), 4% PFA, pH 7.4) for 1-7 days at 4 °C. Following fixation, tissues were dehydrated in 30% sucrose overnight at 4 °C and embedded in OCT. Frozen tissues were sectioned at a thickness of 10 or 50 μm using the cryostat Leica CM1950 and SuperFrost Plus™ Adhesion slides (Epredia).

#### Samples from *M. tuberculosis*-infected guinea pig

FFPE archived samples from previous experiments were kindly provided by Dr. Salguero and Dr. Clark (UK Health Security Agency, United Kingdom). Briefly, samples came from adult female Dunkin-Hartley guinea pigs (*Cavia porcellus*) obtained from a UK Home Office accredited facility (Envigo, United Kingdom). Specific pathogen-free (SPF) animals were individually identified using subcutaneously implanted microchips (Plexx, Netherlands) and housed at ACDP (Advisory Committee on Dangerous Pathogens) level 3, post-infection in groups of up to eight, with access to food and water *ad libitum*. The housing environment was maintained within a temperature range of 15–21 °C and a relative humidity range of 45 to 65%. All animal procedures were approved by the United Kingdom Health Security Agency, Porton Down Establishment Animal Welfare and Ethical Review Body and authorized under an appropriate UK Home Office project license. Animals’ clinical and behavioral status were monitored daily. The *M. tuberculosis* H37Rv strain was used for the infection challenge. National Collection of Type Cultures (NCTC) 7,416 challenge stock was generated from a chemostat grown to steady state under controlled conditions at 37 °C ± 0.1, pH 7.0 ± 0.1 and a dissolved oxygen tension of 10% ± 0.1, in a defined medium, the details of which have been previously described (James et al. 2000). Aliquots were stored at −80 °C. Titer of the stock suspension was determined from thawed aliquots by enumeration of CFUs cultured onto Middlebrook 7H11 OADC selective agar. Animals were challenged using a contained Henderson apparatus in conjunction with an AeroMP control unit as previously described (Hartings and Roy 2004; Clark et al. 2011). Aerosol particles generated were delivered to the animals via aerosol using a 3-jet Collison nebulizer, for an exposure time of 5 min. The challenge suspension in the nebulizer was adjusted by dilution in phosphate buffer saline to a concentration of between 5 × 10^5^ to 1 × 10^6^ CFU/mL to deliver the required estimated, retained, inhaled, dose of 10–20 CFUs to the lungs of each animal. The suspension of *M. tuberculosis* in the nebulizer was plated onto Middlebrook 7H11 OADC selective agar to measure the concentration and confirm retrospectively that the expected doses had been delivered (Clark et al. 2011). Guinea pigs were euthanized at different time-points after infection (day 28 for early-stage lesions, and day 84 for late-stage lesions) by an overdose of sodium pentobarbital via the *i.p.* route, and lung simples were fixed by immersion in neutral buffered formalin (NBF) (Solmedia Ltd., Shrewsbury, United Kingdom) and embedded routinely into paraffin wax (Larenas-Muñoz et al. 2023b).

#### Samples from mice infected with Blastomyces dermatitidis and Histoplasma capsulatum

Lung sections from mice infected with the fungi *Blastomyces dermatitidis* or *Histoplasma capsulatum* were generously provided by Drs. Klein and Wuethrich (University of Wisconsin-Madison, United States) (Kohn et al. 2022). All mouse strains were sourced from the Jackson Laboratory and bred at the specific-pathogen-free BRMS Mouse Breeding Core. Experiments were initiated with mice aged 8 to 12 weeks. Mice were housed and cared for in compliance with the University of Wisconsin Animal Care Committee guidelines, and all experiments followed the IACUC-approved protocol M005891. C57BL6 mice were infected with 20,000 *Blastomyces dermatitidis* yeast or 2 million *Histoplasma capsulatum* yeast by intratracheal aspiration. At 15 days post-infection, mice were sacrificed, and lungs were prepared and embedded in optimal cutting medium (OCT, CellPath) using a protocol similar to the one outlined above.

#### Lung sections from mice infected with SARS-CoV-2

Lung sections from SARS-CoV-2-infected mice were provided by Drs. Montagutelli and Simon-Lorière (Institut Pasteur, France). All mouse experiments were conducted in compliance with French and European regulations (Directive 2010/63EU) and received approval from the Institut Pasteur Ethics Committee (Protocol dap210050), authorized by the French Ministry of Research (authorization 31816). B6.Cg-Tg(K18-ACE2)2Prlmn/J transgenic mice were purchased from The Jackson Laboratory and bred at the Institut Pasteur under SPF conditions. Infection was carried out in isolators, within the BSL-3 mouse facility at the Institut Pasteur. Mice were fed standard chow *ad libitum* and kept in a 14:10 light:dark cycle. The D614G variant (B.1) strain was supplied by the National Reference Centre for Respiratory Viruses hosted by Institut Pasteur (Paris, France), led by Pr. Sylvie van der Werf, through the European Virus Archive goes Global (Evag) platform funded by the European Union’s Horizon 2020 research and innovation program (grant 653316). Mice were anesthetized (ketamine/xylazine) and inoculated *i.n.* with 4.5×10^4^ plaque-forming units (PFU) of SARS-CoV-2 in a 40 μL PBS. At day 6 post-infection, mice were euthanized by ketamine/xylazine overdose. The left lung lobe was fixed in 10% phosphate-buffered formalin for 24-36 hours, transferred into 70% ethanol and processed for histological analysis using paraffin-embedded, 4 μm-thick sections.

#### Samples from asthma model in mice

Samples came from Dr. Gaudenzio (Infinity, France). Isoflurane-anesthetized C57BL/6 mice were instilled *i.n*. with 100 ng of lipopolysaccharide (LPS, Sigma-Aldrich). Vehicle mice were instilled *i.n.* with 50 µL of PBS. One day later (day 1), mice were administered *i.n.* with 40 µg of house dust mites (HDM) pteronyssinus (Greer Laboratories) in 50 µL of PBS. Seven days later (day 8), all mice were challenged by *i.n.* instillation of 10 µg of HDM in 50 µl of PBS. Three days after the HDM challenge (day 11), animals were euthanized, and lungs were embedded in OCT using a protocol similar to the one outlined above.

#### Tissue preparation for immunohistofluorescence

Paraffin-embedded samples were first processed by Heat-Induced Epitope Retrieval (HIER) with 10 mM citrate buffer, pH6, to retrieve antigens before the blocking step, while OCT-embedded samples were thaw in milli-Ro water for 30 minutes at room temperature. All sections were blocked in PBS containing 5% goat serum (GeneTex), 5% bovine serum albumin (BSA, Euromedex) and 0.2% Triton X100 (Sigma-Aldrich) for three hours at room temperature. Tissues were then incubated overnight with primary antibodies (listed in the figure legend and (**Table S3**) diluted in PBS with 5% BSA and 0.2% Triton X100. Appropriate isotype controls were included. After washing, the sections were stained with secondary antibodies (**Table S3**) for four hours at RT, followed by DAPI (Sigma-Aldrich) staining for ten minutes at room temperature. Sections were then mounted with DAKO fluorescent mounting medium (Agilent Technologies) and images were acquired using a Zeiss AXIO Imager-M2 epifluorescence microscope and a Zeiss LSM 710 confocal microscope. The images were processed in ImageJ using the *Despeckle* filter to attenuate or remove bright, isolated pixels. For 2D imaging, the workflow continued in ImageJ with the application of *z-projection*. A *Median* filter with a 2-pixel radius was subsequently applied to further enhance image quality. When necessary, local gamma correction was selectively applied to the DAPI staining to optimize tissue visualization. In human lung biopsies presenting high levels of fibrosis, *Image calculator* was used to subtract non-specific double positive signals between green and red channels an allow the interpretation of the images. For 3D reconstructions and movie generation, the images were processed using Imaris software.

#### Quantification of inflammation and TUBB3^+^ surfaces

The software QuPath was used to quantify inflammation and TUBB3-positive signal in lung sections from *M. tuberculosis*-infected mice. Bronchi and inflamed areas/lesions were delimitated, using *Polygon* tool, based on morphological criteria. Inflamed and TUBB3^+^ areas were segmented, and a threshold was applied to the image to distinguish between positive and negative signals using *Classify / Pixel classification / Threshold*. The parameters were kept constant throughout the quantification process.

#### Quantification of distance between cells

The software Imaris was used to quantify the distance between cells, or between cells and bacteria. TUBB3 staining was converted into a surface, followed by the application of the *Distance transform of the outside surface object* tool to create a gradient of distances around each cell. Next, immune cells or bacteria were also converted into surfaces, and the *Intensity mean* statistics was used to determine their distance within the gradient.

#### Tissue clearing

The polyethylene glycol (PEG)-associated solvent system (PEGASOS) protocol was used for tissue clearing of lungs (Jing et al. 2018). Briefly, both non-infected and *M. tuberculosis*-infected mice were euthanized, and a transcardiac perfusion with 50 mL of PBS was performed to remove blood. The lungs were then harvested and fixed in a 4% PFA for 24 hours. Following fixation, samples were treated with 25% Quadrol decolorization solution (Sigma-Aldrich) at 37 °C for two days. Next, samples were immersed in gradient delipidation solutions (30%, 50%, 70% tert-butanol (Sigma-Aldrich), and 3% Quadrol) at 37 °C shaker for one to two days. For the whole-mount immunohistofluorescence staining, samples were incubated overnight at room temperature in blocking buffer containing 10% DMSO (Sigma-Aldrich), 0.5% IgeePal630 (Sigma-Aldrich), 10% goat serum and PBS. Blocking step was followed by incubation with primary antibody incubation (TUBB3, see **Table S3**) in blocking buffer at 4 °C shaker for three days, washing in PBS for one day shaker, and incubation with secondary antibody (see **Table S3**) in blocking buffer at 4 °C shaker for two days. Next, samples were washed in PBS for six hours shaker before to be treated with dehydration solution (70% tert-butanol / 30% PEG-MMA-500 (Sigma-Aldrich) / 3% Quadrol) for an additional one to two days. The clearing process was completed by immersing the samples in BB-PEG clearing medium (70% benzyl benzoate (Sigma-Aldrich) / 27% PEG-MMA-500 / 3% Quadrol) for at least one day until reaching transparency. Finally, the cleared samples were preserved in the clearing medium at room temperature. Imaging was performed by the company Imactiv-3D using light-sheet microscopy within a week.

#### Image stack cleaning

To proceed with the 3D reconstruction, non-specific signals from secondary antibody had to be removed. The entire procedure was carried out on a computer workstation (Dell Precision Intel Xeon 3.9 GHz, 256 Go RAM) and with a standard installation of Fiji. First, bright isolated pixels in the image stack were attenuated or even removed by applying 20 times the *Despeckle* filter. Next, fluorescent grain detection was performed using *Weka segmentation* plugin. The model was trained with two classes (class 1 grain, class 2 background) until optimal segmentation was achieved, with pre-processing playing a crucial role in isolating grains. The resulting segmentation was converted to a mask, and grains were detected and stored in the ROI Manager using the *Analyze Particles* function. These ROIs were then applied to the original image stack, and *Smoothing* function was used to reduce graininess, repeating the process 100 times. Once the images were sufficiently refined, 3D reconstruction and movie generation were completed using Imaris software.

#### Statistical analysis

Statistical information, including n, statistical tests, and statistical significance values, is indicated in the figure legends and analysis was performed using Prism 10.0 (GraphPad). p<0.05 was considered as the level of statistical significance (*p<0.05; **p<0.005; ***p<0.0005; ****p<0.0001).

## Acknowledgments

We thank Céline Beronne, Flavie Moreau, Malory Blasco, Aline Tridon and Amaury Schneider (Genotoul Anexplo-IPBS platform, Toulouse, France, member of the Celphedia national infrastructure) for mouse care, and Emmanuelle Näser (Genotoul TRI-IPBS platform, Toulouse, France, member of the France-BioImaging national infrastructure supported by the French National Research Agency, ANR-10-INBS-0005 FBI BIOGEN) for assistance with flow cytometry. We also acknowledge Denis Hudrisier for the advice provided and sharing the material related to the lungs of *M. tuberculosis*-infected *Sp140*^-/-^ mice. We thank the histological IPBS platform for helping with sample preparation and analysis. We acknowledge the help of Elizabeth Bellard and Eve Pitot with imaging. We thank Jean-Michel Lagarde, Aurélie Gomes and Pascale Bernes-Lasserre (Imactiv-3D, Toulouse, France) for 3D reconstructions using light-sheet microscopy. We also acknowledge Dr. Jean-Philippe Girard for sharing his microscope. We thank Dr. Lillian Basso (Infinity, Toulouse, France), Dr. Mauro Gaya (CIML, Marseille, France) and Dr. Russel Vance (UC Berkeley) for providing mice. Dr. Marcelo J. Kuroda and Dr. Deepak Kaushal for providing the NHP samples from the study supported by NIH award OD011104, AI111943, AI111914, AI097059 and AI110163. We extend our gratitude to Laurine Conquet and Matthieu Prot for their technical support, as well as the staff of the BSL-3 mouse facility and the histology platform, both part of the C2RA at the Institut Pasteur. We thank Prof. Philippe Manivet, head of the Centre de Ressources Biologiques (CRB-LRB), and Prof. Philippe Bertheau, Head of the Department of Anatomic Pathology at Hôpital Saint-Louis, for facilitating the collection of lymph node biopsies from tuberculosis-infected patients. This project was financially supported by the Centre National de la Recherche Scientifique, Université Paul Sabatier, ANRS Maladies Infectieuses Émergentes (Grants ECTZ242543, ECTZ 2093306 to O.N. and C.V.), the Fondation pour la Recherche Médicale (Grant EQU202103012733 to O.N.), and the Fondation Bettencourt Schueller (Grant Explore-TB to O.N.). The acquisition of lung sections from SARS-CoV-2-infected mice was supported by the French Government’s Investissement d’Avenir program, Laboratoire d’Excellence Integrative Biology of Emerging Infectious Diseases (Grant No. ANR-10-LABX-62-IBEID). The acquisition of lung tissue from fungus-infected mice was supported by the grants 01AI040996 (BK/MW), R01 AI168370 (BK), U01 AI124299 (BK), R37 AI035681 (BK), R01 AI168370 (BK). The clinical study SH-TBL, from which the human lung samples were obtained, was funded by the Plan Nacional I + D + I co-financed by ISCIII-Subdirección General de Evaluación and Fondo-EU de Desarrollo Regional (FEDER) (PI16/01511, PI20/01424, CP13/00174, CPII18/00031 and CB06/06/0031); the Catalan Agency for Management of University and Research Grants (AGAUR) (2021 SGR 00920); and the “Spanish Society of Pneumology and Thoracic Surgery” (SEPAR) (16/023).

## Supplemental Figure legends

**Supplemental Figure 1. TUBB3^+^ cells accumulate in lung lesions of *M. tuberculosis*-infected mice.**

**A.** Representative immunohistofluorescence images of lungs from non-infected (left) and *M. tuberculosis*-infected (middle, right, H37Rv-dsRed strain in orange) mice at day 42 post-infection, after tissue clearing, showing TUBB3 (green) staining. Scale bar, 300 µm. **B.** Representative immunohistofluorescence images of murine lungs during a time-course of *M. tuberculosis* infection (H37Rv-dsRed strain in orange). TUBB3 (green) and nuclei (DAPI, blue) are shown. Data represent three independent experiments, with n=4 mice per timepoint. Scale bar, 500 µm. **C.** Percentage of TUBB3-stained surface per lobe of *M. tuberculosis*-infected murine lungs. Kruskal-Wallis test with Dunn’s multiple group comparisons, three independent experiments, n=4 mice per timepoint. **D.** Percentage of inflammation surface per lobe of *M. tuberculosis*-infected murine lungs, defined by areas with high numbers of nuclei. Kruskal-Wallis test with Dunn’s multiple group comparisons, three independent experiments, n=4 mice per timepoint. **E.** Percentage of TUBB3-stained surface in non-inflamed (open circles) and inflamed (filled circles) areas within the same lobe of *M. tuberculosis*-infected lungs. Two-way ANOVA with Sidak’s multiple comparisons, three independent experiments, n=4 mice per timepoint. **F.** Quantification of the distance between bacteria and TUBB3^+^ cells from at least three mice across 2-3 independent experiments. *, p≤0.05; **, p≤0.01; ***, p≤0.001; ****, p≤0.0001.

**Supplemental Figure 2. TUBB3^+^ cells do not express markers for epithelial cells, endothelial cells, fibroblasts, muscle cells, or immune cells.**

**A-E.** Representative immunohistofluorescence images (left) of lungs from mice infected with *M. tuberculosis* H37Rv-dsRed strain (orange) at day 42 post-infection. TUBB3 (green) is shown alongside markers for epithelial cells (E-Cadherin, magenta, A), endothelial cells (CD31, magenta, B), fibroblasts (Vimentin, magenta, C), muscle cells (α-SMA, magenta, D), and immune cells (CD45, magenta, E). Histograms (right) depict the fluorescence intensity along the dotted lines from the images on the left. Data represent two independent experiments, with n=3 mice. Scale bar, 20 µm.

**Supplemental Figure 3. TUBB3^+^ cells do not express markers for nociceptors, sympathetic nerves, or parasympathetic nerves.**

**A-B.** Representative immunohistofluorescence images of lungs from *M. tuberculosis* (H37Rv-dsRed strain, orange)-infected mice at day 42 post-infection. TUBB3 (green), TH (magenta, B), VAChT (magenta, C) and nuclei (DAPI, blue) are shown. Images 1 show zoomed views of bronchi (dotted lines), and images 2 show zoomed views of the parenchyma. Data represent two independent experiments, with n=6 mice. Scale bar, 100 µm. **C.** Representative immunohistofluorescence images of lungs from TRPV1-YFP (magenta) mice infected with *M. tuberculosis* H37Rv-dsRed strain (orange) for 28 days. TUBB3 (green) and nuclei (DAPI, blue) are shown. Images 1 show zoomed views of bronchi (dotted lines), and images 2 show zoomed views of the parenchyma. Data represent two independent experiments, with n=6 mice. Scale bar, 100 µm.

**Supplemental Figure 4. A. The presence of TUBB3^+^ cells is not affected by the depletion of Na_V_1.8^+^ nociceptors.** Representative immunohistofluorescence images of lungs from *M. tuberculosis* (H37Rv-dsRed strain, orange)-infected control DTA and Na_V_1.8-DTA mice at day 28 post-infection. TUBB3 (green) and nuclei (DAPI, blue) are shown. n=3 mice per group. Scale bar, 500 µm. **B-E. The emergence of TUBB3^+^ cells coincide with the recruitment of DCX^+^ PSA-NCAM^+^ neuronal progenitor-like cells.** Representative immunohistofluorescence images of lungs from *M. tuberculosis*-infected mice at day 42 post-infection. DCX (magenta) and nuclei (DAPI, blue) are shown along with PSA-NCAM (yellow, C), Ki-67 (yellow, D) and CD45 (yellow, E). Data represent two independent experiments, with n=3 mice. Scale bar, 50 µm.

**Supplemental Figure 5. TUBB3^+^ cells colonize *M. tuberculosis*-induced lesions and iBALT, interact with CD4^+^ and CD8^+^ T cells, but are unlikely to present antigens.**

**A.** Representative immunohistofluorescence images showing the proximity of TUBB3^+^ cells (green) to macrophages (F4/80, yellow) and T cells (CD3, magenta) in *M. tuberculosis*-induced lesions in murine lungs at day 42 post-infection. Nuclei are stained with DAPI (blue). Data represent 2-3 independent experiments, with n=3 mice. Scale bar, 100 µm. **B.** Representative immunohistofluorescence images showing an iBALT in *M. tuberculosis*-infected murine lungs at day 42 post-infection. TUBB3 (green), nuclei (DAPI, blue), follicular dendritic cells (CD21/35, magenta) and B cell (B220, cyan) are shown. Data represent two independent experiments with n=3 mice. Scale bar, 50 µm. **C.** Representative immunohistofluorescence images showing the proximity of TUBB3^+^ cells (green) to CD4^+^ T cells (magenta, left) or CD8^+^ T cells (magenta, right) in *M. tuberculosis*-infected murine lungs at day 42 post-infection. Nuclei are stained with DAPI (blue). Data represent two independent experiments, with n=3 mice. **D.** Quantification of the distance between CD4^+^ and CD8^+^ T cells and TUBB3^+^ cells, based on images in C. **E-F.** Representative immunohistofluorescence images (left) of *M. tuberculosis*-infected murine lungs at day 42 post-infection. TUBB3 (green), MHC-I (magenta, E) and MHC-II (magenta, F) are shown. Nuclei are stained with DAPI (blue). Histograms (right) depict the fluorescence intensity along the dotted lines from the corresponding images on the left. Scale bar, 20 µm.

**Supplemental Figure 6. IL-17A is dispensable for the recruitment of TUBB3^+^ cells upon TB infection.**

Representative immunohistofluorescence images of *M. tuberculosis*-infected lungs of control (left) and *Il17a*^-/-^ (right) mice. TUBB3 (green), CD3 (T cells, magenta), F4/80 (macrophages, yellow) and nuclei (DAPI, blue) are shown. Parenchymal TUBB3^+^ cells are indicated with white arrowheads in the zoomed lower panels. n=3 mice/conditions. Scale bar, 200 μm.

**Supplemental Figure 7. TUBB3^+^ cells are found in the lung of *M. tuberculosis*-infected guinea pigs.**

Representative immunohistofluorescence images of lungs from *M. tuberculosis*-infected guinea pigs at day 28 and 84 post-infection. TUBB3 (green), nuclei (DAPI, blue), CD68 (macrophages, yellow) and CD3 (T cells, magenta) are shown. Data represent n=1 animal per timepoint. Scale bar, 100 µm.

**Supplemental Figure 8. TUBB3^+^ cells are observed in lung and lymph node biopsies of TB patients.**

**A.** Representative immunohistofluorescence images of TUBB3^+^ area from lung biopsies of TB patient, (Patient TB031, left; TB018 middle; TB013, right, see **Table S1**). TUBB3 (green), nuclei (DAPI, blue), CD68 (macrophages, yellow) and CD3 (T cells, magenta) are shown. Data represent n=11 patients analyzed with similar features. White arrowheads indicate parenchymal TUBB3^+^ cells. Scale bar, 200 µm. **B.** Number of TUBB3^+^ areas in lung biopsies from non-TB patients (n=2) and TB patients (n=11). Mean values are represented. Mann-Whitney test, *, p ≤ 0.05. **C-E.** Number of TUBB3^+^ lesions in (**C**) drug-sensitive (DS-TB, n=4) *vs*. multidrug-resistant (MDR-TB, n=7) TB patients, (**FD**) newly diagnosed TB patients (n=6) *vs*. patients with relapse (n=4), and (**GE**) male (n=7) *vs*. female (n=4) TB patients. Box and whisker plots (min to max) display medians as horizontal lines. **F.** Representative immunohistofluorescence images of lymph node biopsies from TB patients. TUBB3 (green), CD68 (macrophages, yellow) and CD3 (T cells, magenta) are shown for two patients (Patients 63924 and 51102; see **Table S2**). Data represent n=6 patients analyzed with similar features. White arrowheads indicate parenchymal TUBB3^+^ cells. Scale bar, 100 µm.

**Movie 1:**
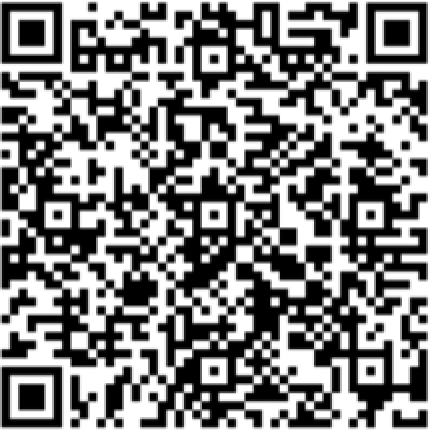
Tissue clearing and 3D reconstruction of non-infected and *M. tuberculosis*-infected mouse lungs stained for TUBB3. TUBB3 (green) and bacilli (*M. tuberculosis*, orange) are shown. Scale bar, 300 µm for videos 1 and 2, and 100 µm for video 3.

**Movie 2:**
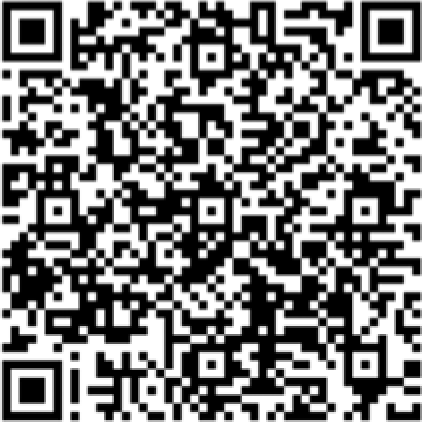
Morphology of TUBB3^+^ cells. TUBB3 (green), bacilli (*M. tuberculosis*, orange) and nuclei (DAPI, blue) are shown. Scale bar, 5 µm.

**Movie 3:**
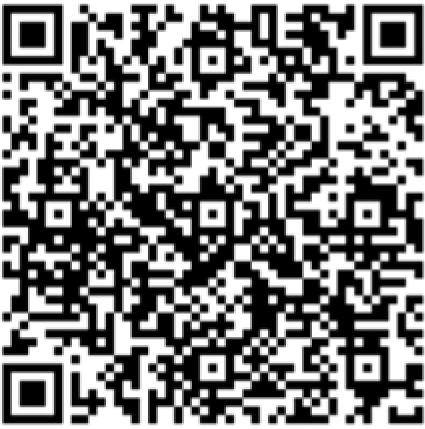
TUBB3^+^ cells are not in a proliferative state. TUBB3 (green), Ki67 (magenta) and nuclei (DAPI, blue) are shown. Scale bar, 5 µm.

**Movie 4:**
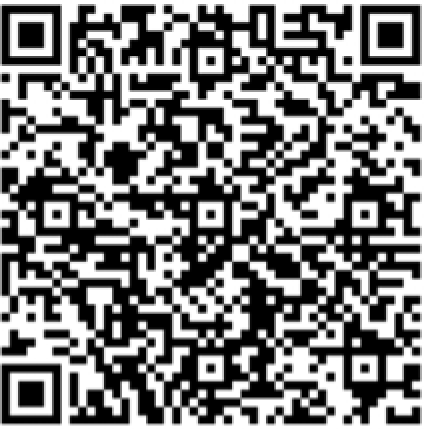
TUBB3 cells are in close contact with immune cells. TUBB3 (green), macrophages (F4/80, yellow), B cells (B220, cyan), T cells (CD3, CD4, CD8, magenta), and nuclei (DAPI, blue) are shown. Scale bar, 5 µm.

## SUPPLEMENTARY TABLES

**Supplementary File - Table S1.**
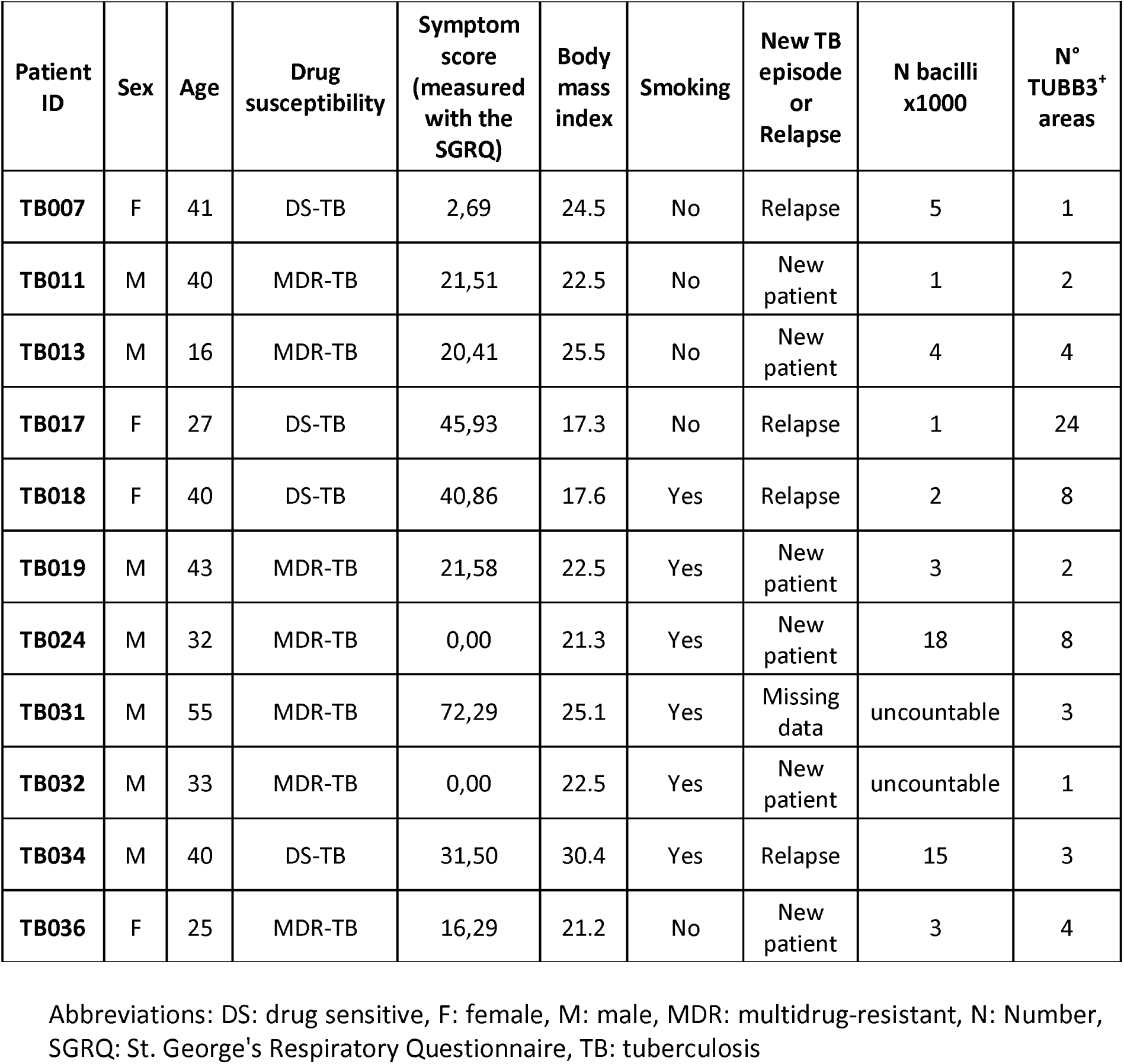
Clinical data of TB patients in the lung biopsy cohort.

| Patient ID | Sex | Age | Drug susceptibility | Symptom score (measured with the SGRQ) | Body mass index | Smoking | New TB episode or Relapse | N bacilli x1000 | N° TUBB3+ areas |
| --- | --- | --- | --- | --- | --- | --- | --- | --- | --- |
| TB007 | F | 41 | DS-TB | 2,69 | 24.5 | No | Relapse | 5 | 1 |
| TB011 | M | 40 | MDR-TB | 21,51 | 22.5 | No | New patient | 1 | 2 |
| TB013 | M | 16 | MDR-TB | 20,41 | 25.5 | No | New patient | 4 | 4 |
| TB017 | F | 27 | DS-TB | 45,93 | 17.3 | No | Relapse | 1 | 24 |
| TB018 | F | 40 | DS-TB | 40,86 | 17.6 | Yes | Relapse | 2 | 8 |
| TB019 | M | 43 | MDR-TB | 21,58 | 22.5 | Yes | New patient | 3 | 2 |
| TB024 | M | 32 | MDR-TB | 0,00 | 21.3 | Yes | New patient | 18 | 8 |
| TB031 | M | 55 | MDR-TB | 72,29 | 25.1 | Yes | Missing data | uncountable | 3 |
| TB032 | M | 33 | MDR-TB | 0,00 | 22.5 | Yes | New patient | uncountable | 1 |
| TB034 | M | 40 | DS-TB | 31,50 | 30.4 | Yes | Relapse | 15 | 3 |
| TB036 | F | 25 | MDR-TB | 16,29 | 21.2 | No | New patient | 3 | 4 |

Abbreviations: DS: drug sensitive, F: female, M: male, MDR: multidrug-resistant, N: Number, SGRQ: St. George's Respiratory Questionnaire, TB: tuberculosis

**Supplementary File - Table S2.**
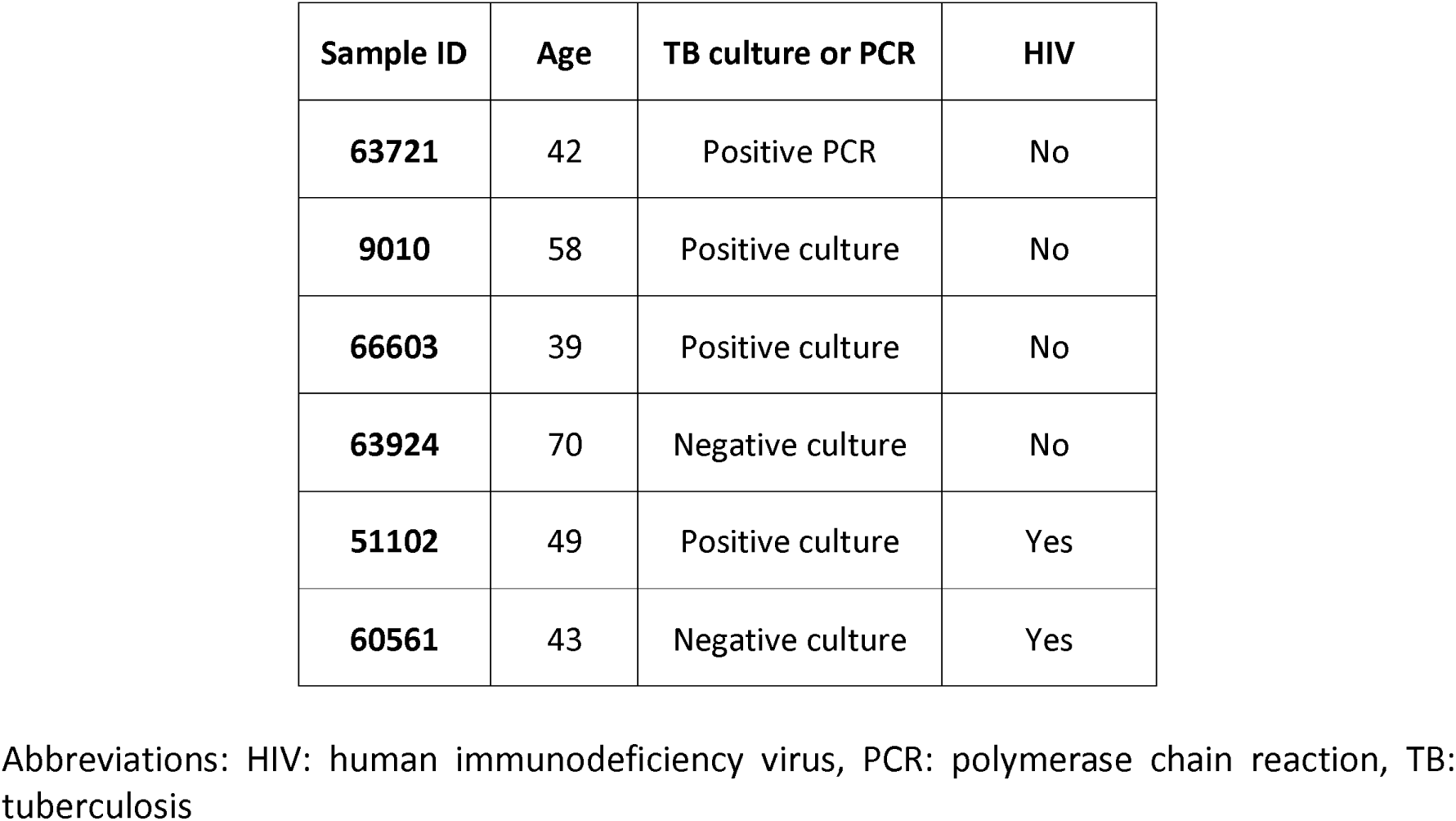
Clinical data of TB patients in the lymph node biopsy cohort.

| **Sample ID** | **Age** | **TB culture or PCR** | **HIV** |
| --- | --- | --- | --- |
| **63721** | 42 | Positive PCR | No |
| **9010** | 58 | Positive culture | No |
| **66603** | 39 | Positive culture | No |
| **63924** | 70 | Negative culture | No |
| **51102** | 49 | Positive culture | Yes |
| **60561** | 43 | Negative culture | Yes |

Abbreviations: HIV: human immunodeficiency virus, PCR: polymerase chain reaction, TB: tuberculosis

**Supplementary File - Table S3.**
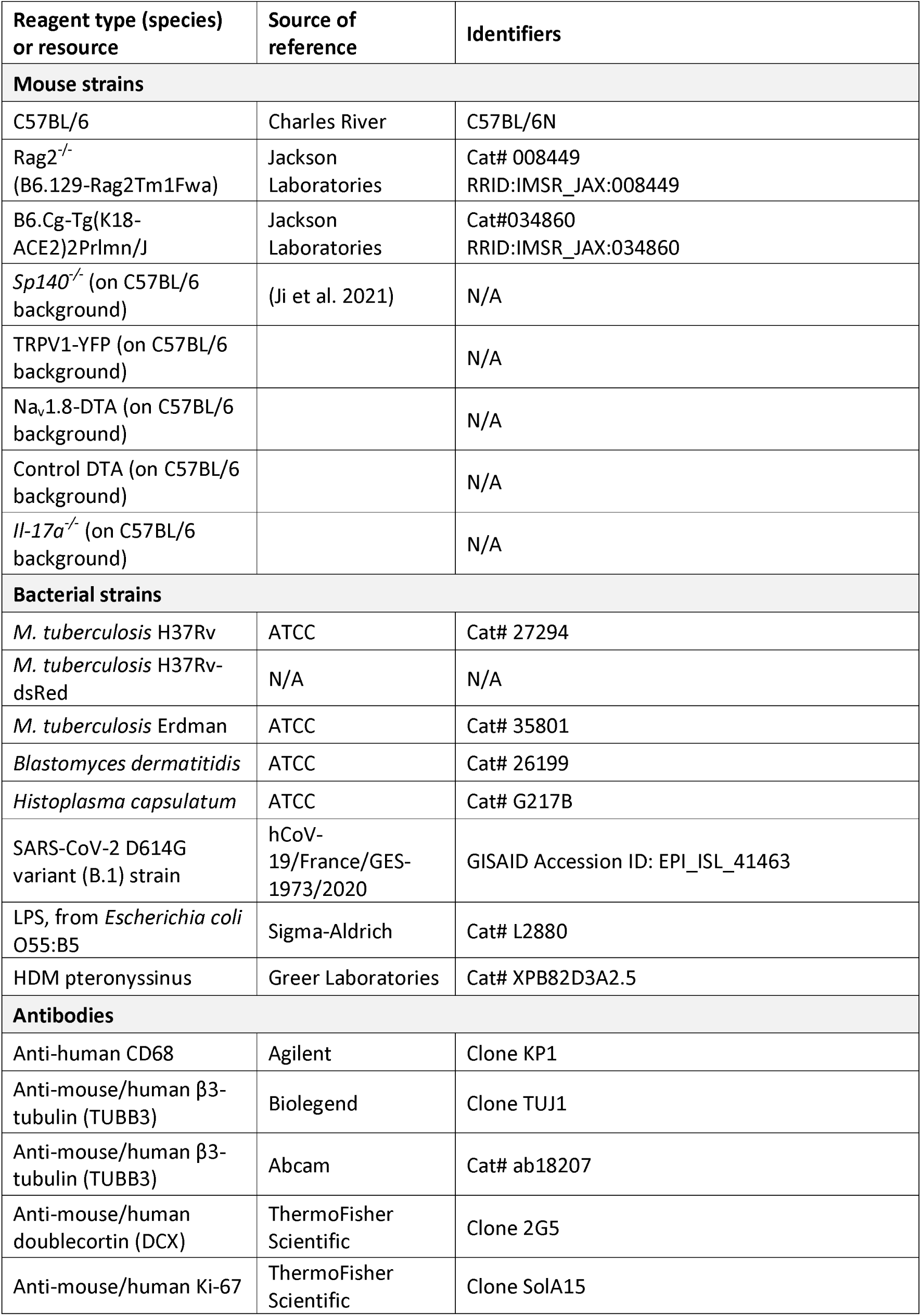
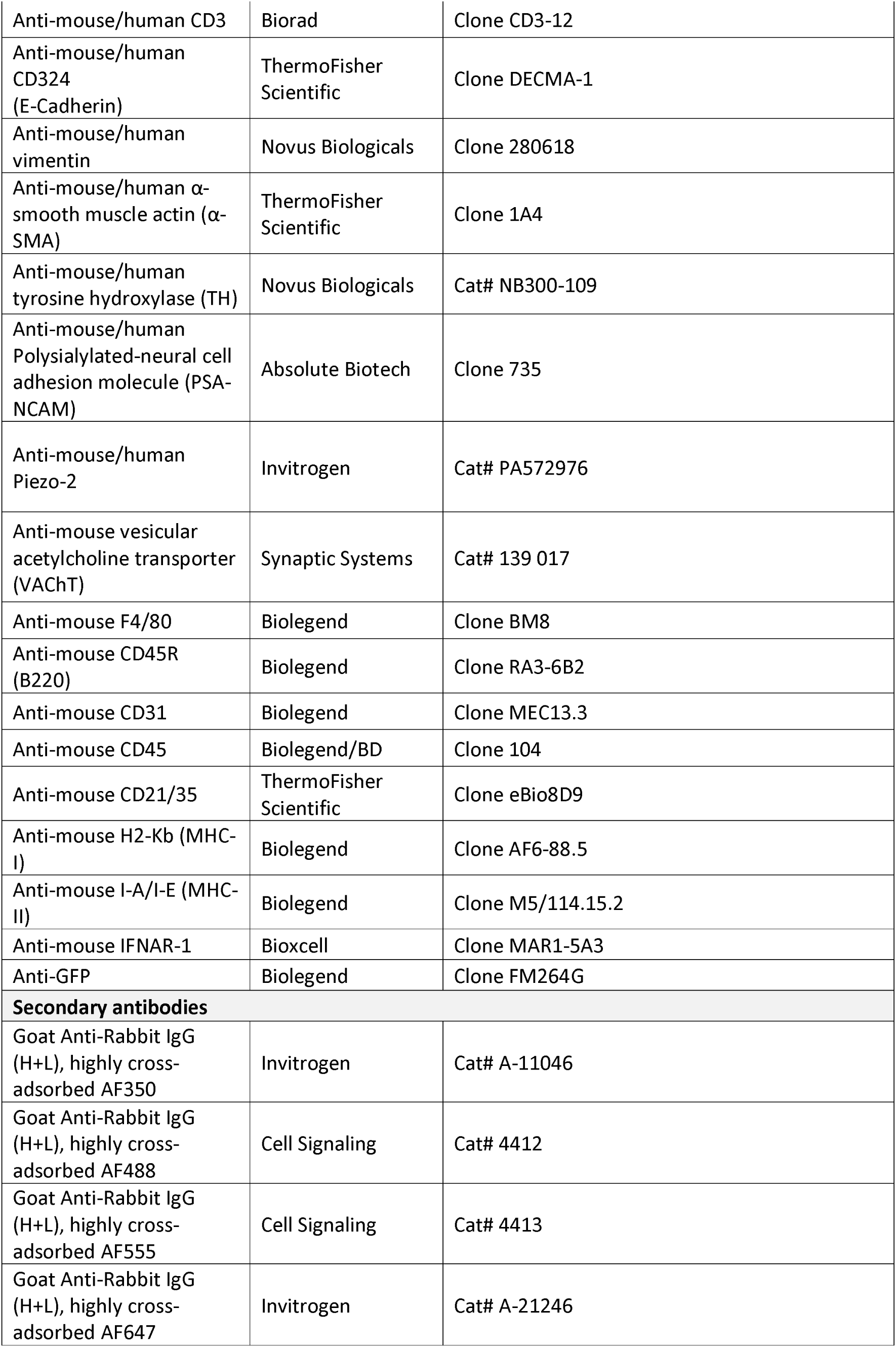
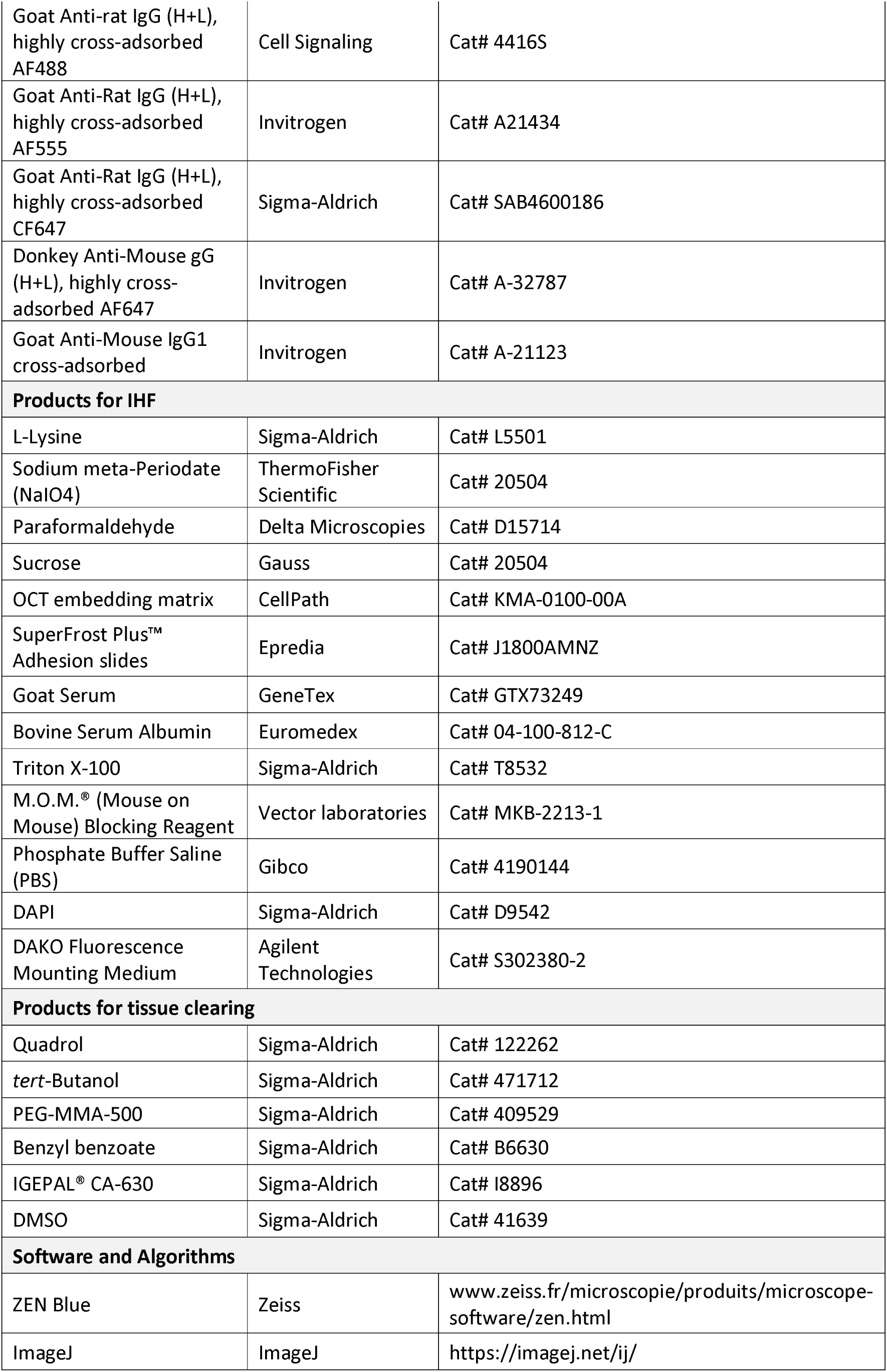
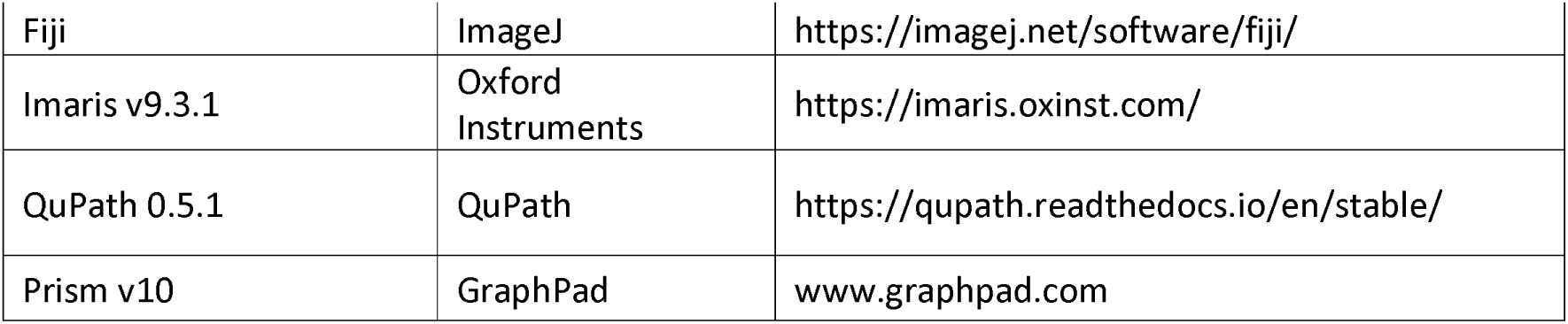
Star Method.

