## Supplementary figures and images for "Identification of a unique TUBB3^+^ cell population in the tuberculosis granuloma"

### Supplemental Figure 1

**Monard, et al. Supplemental Figure 1**

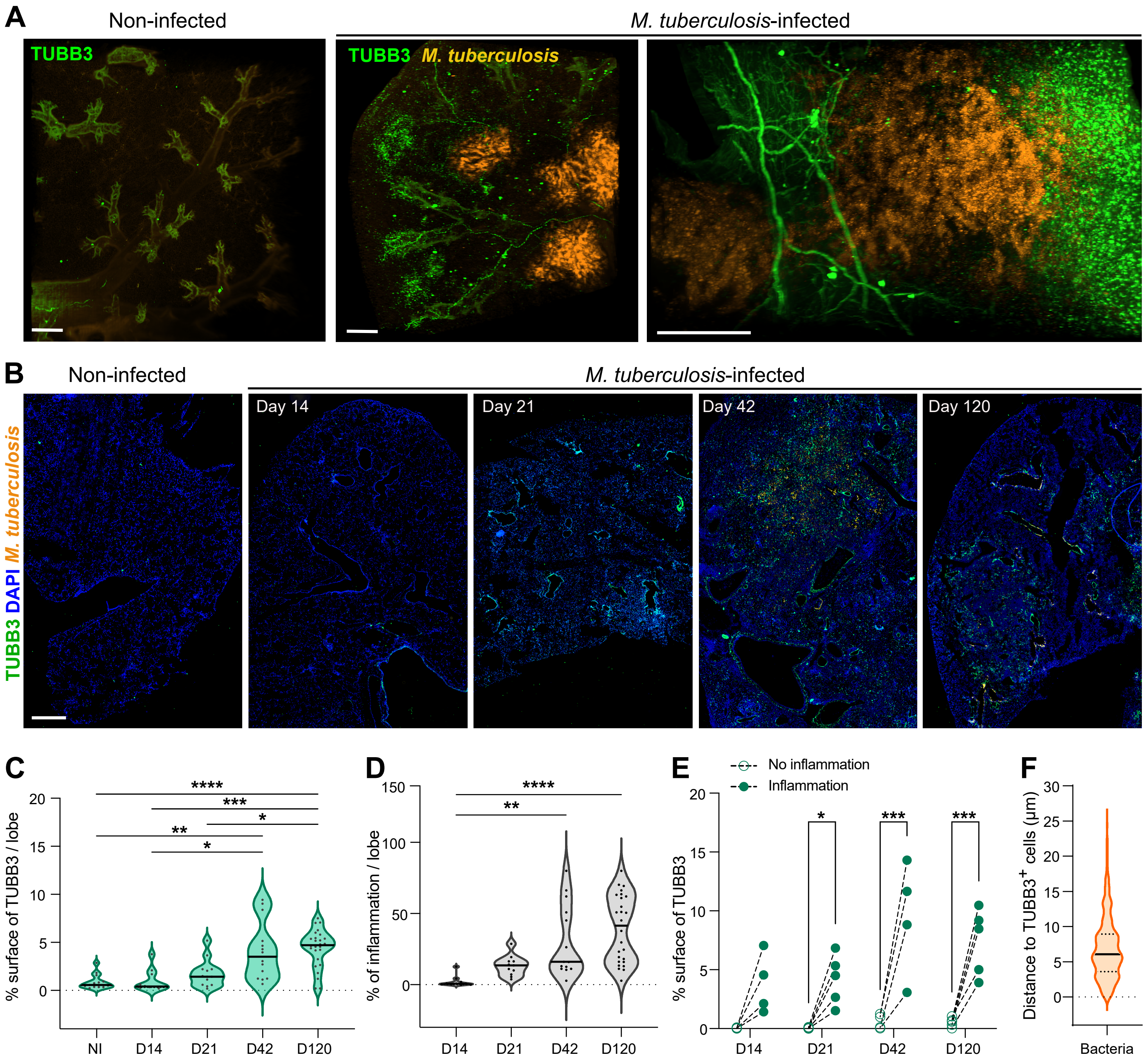

### Supplemental Figure 2

#### Monard et al. Supplemental Figure 2

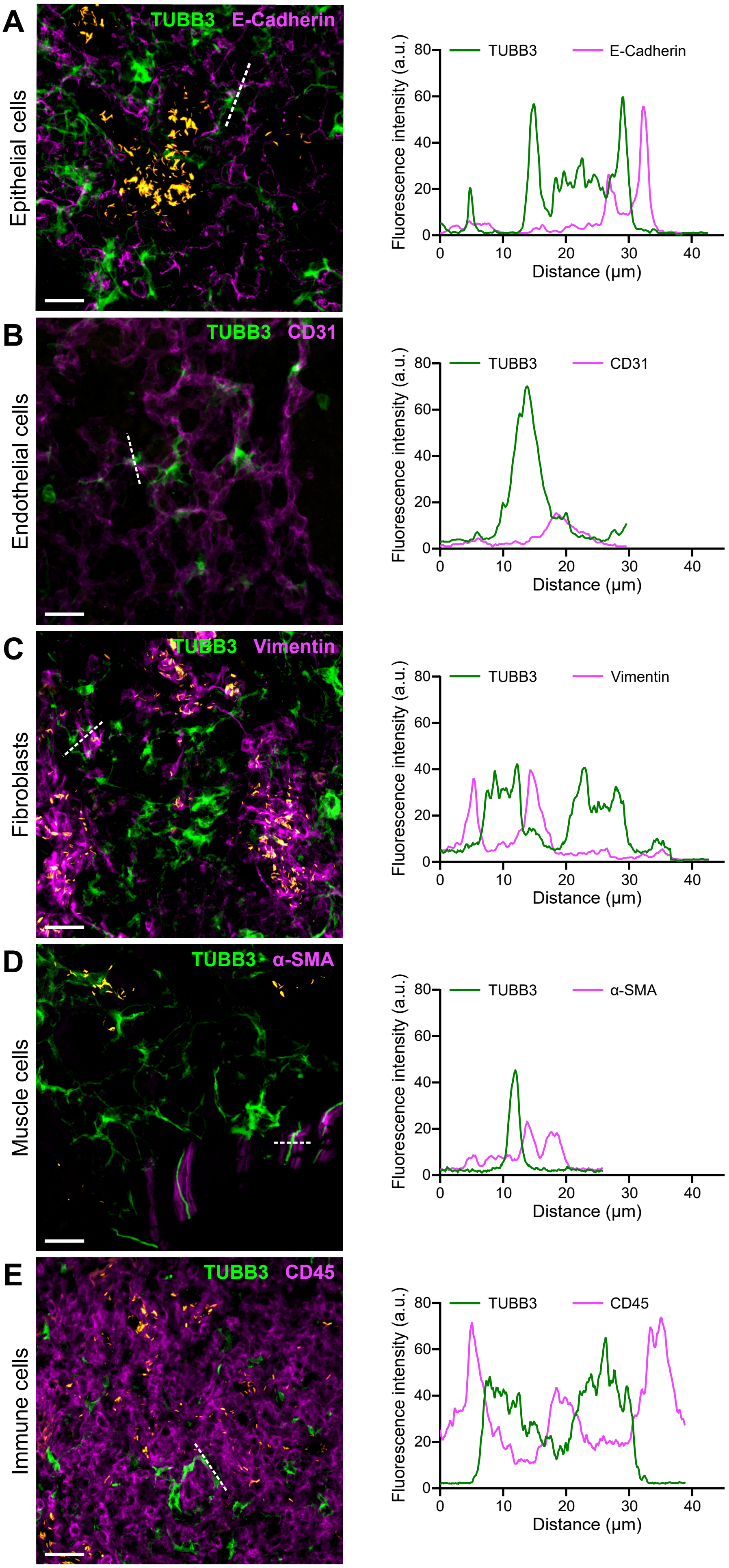

### Supplemental Figure 3

##### Monard, et al. Supplemental Figure 3

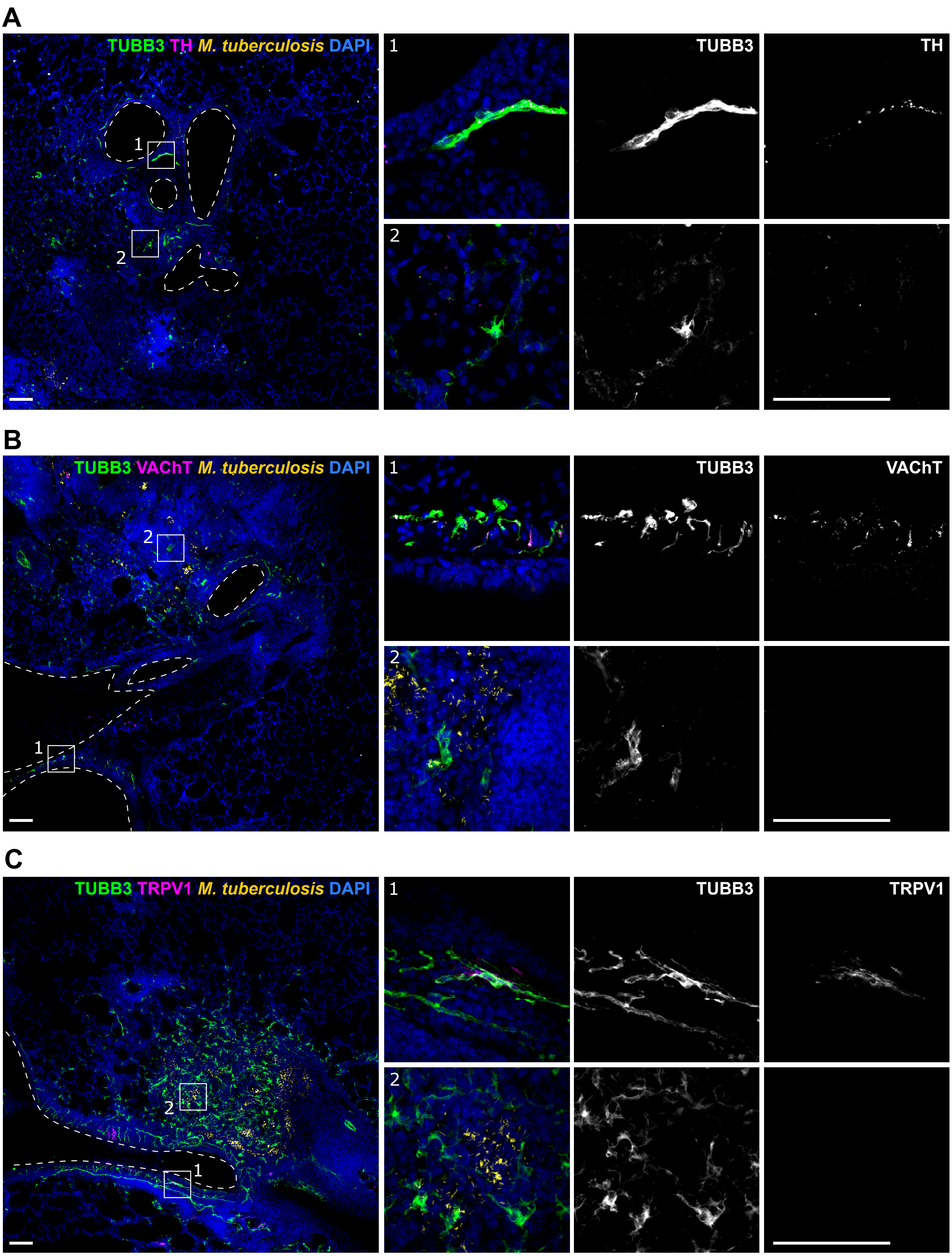

### Supplemental Figure 4

**Monard, et al. Supplemental Figure 4**

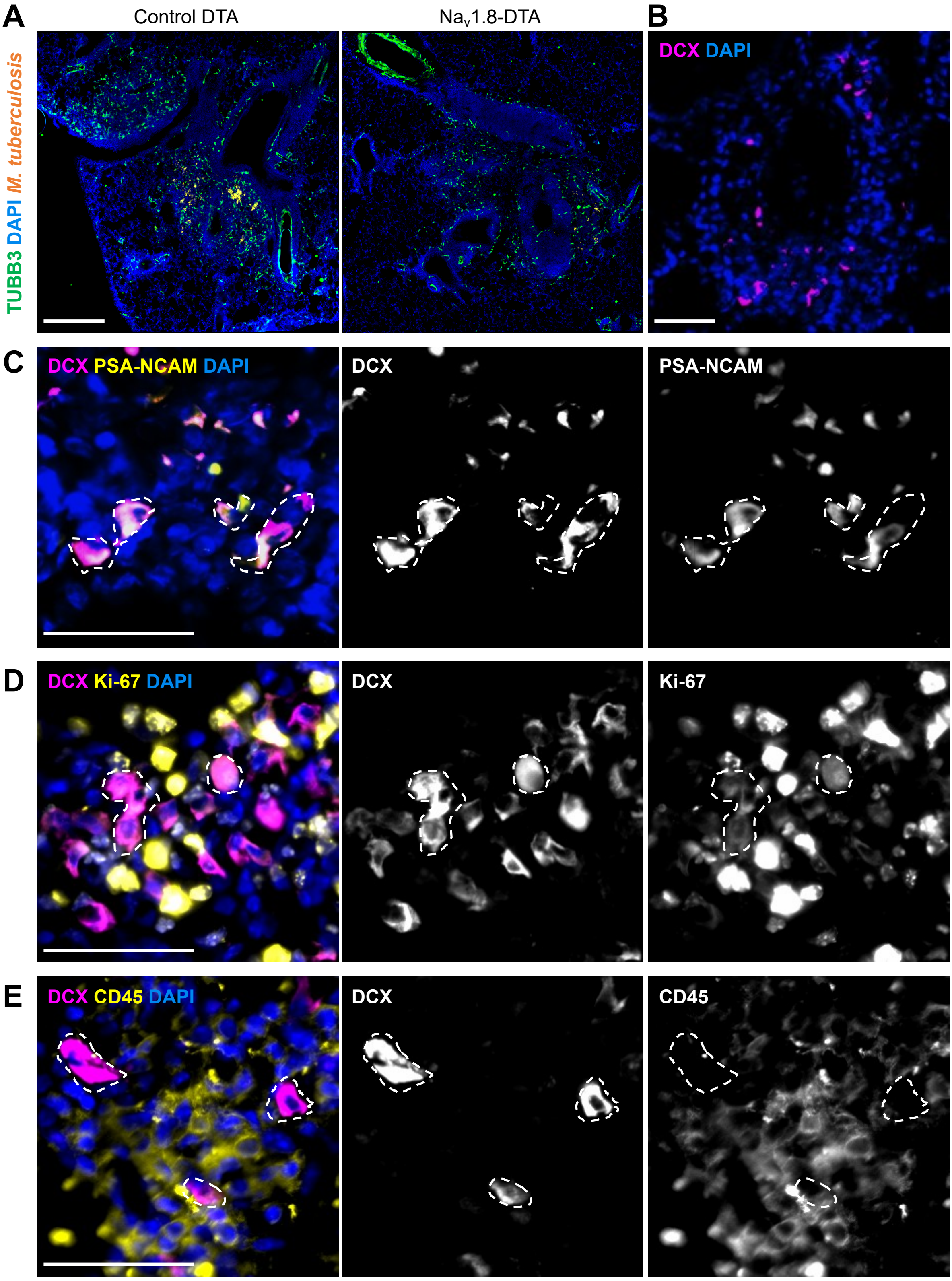

### Supplemental Figure 5

#### Monard, et al. Supplemental Figure 5

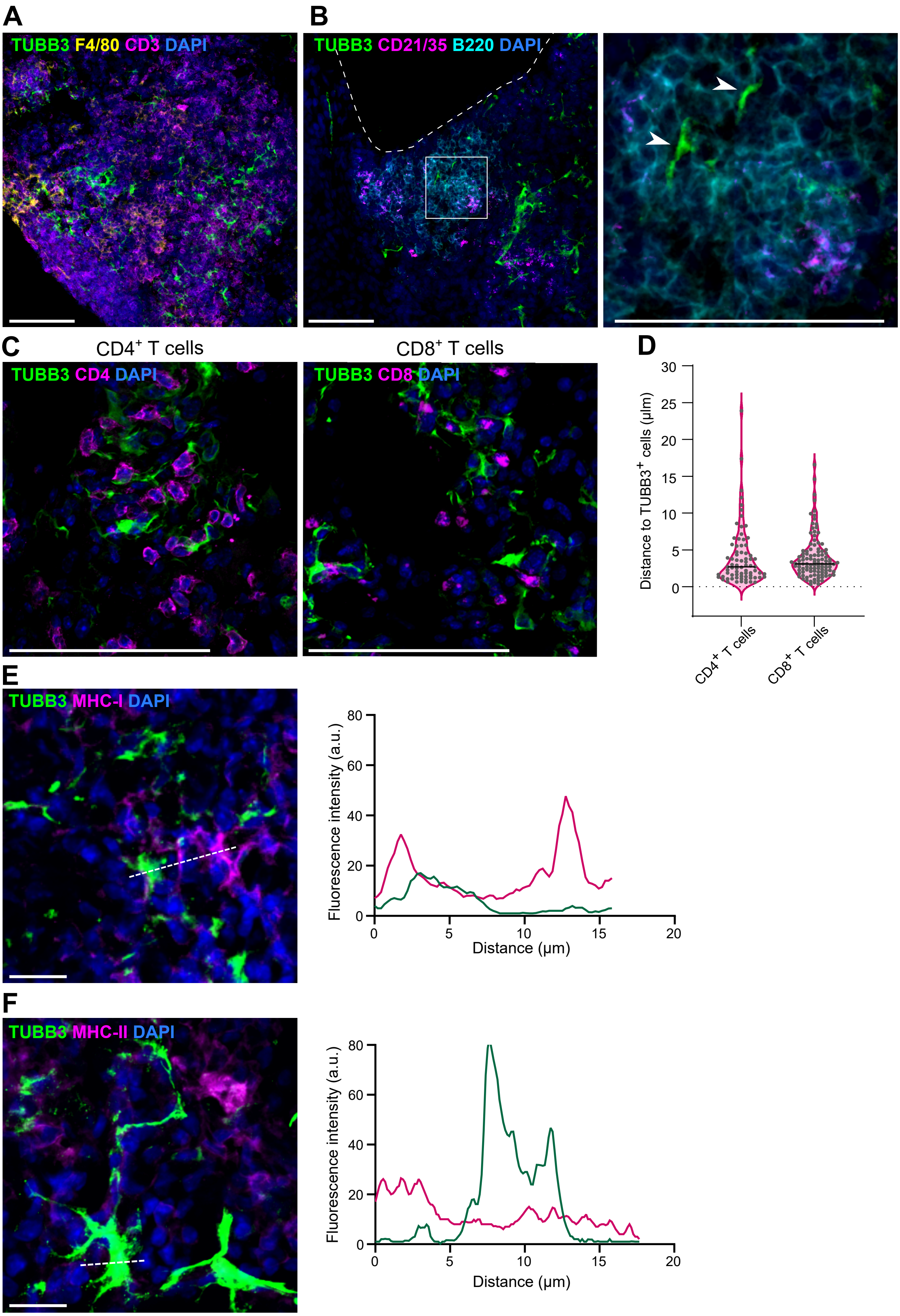

### Supplemental Figure 6

#### Monard et al., Supplemental Figure 6

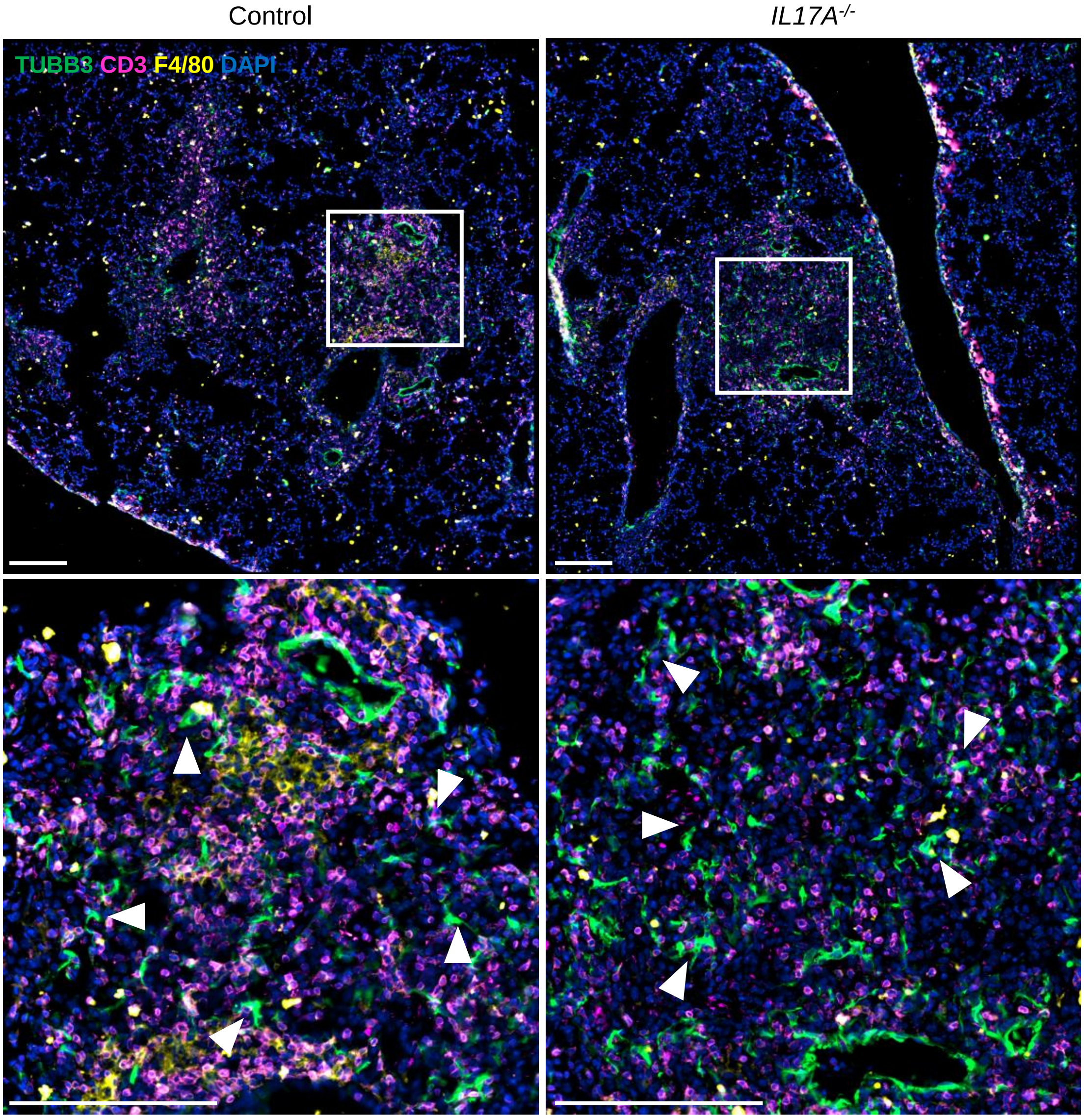

### Supplemental Figure 7

#### Monard, et al. Supplemental Figure 7

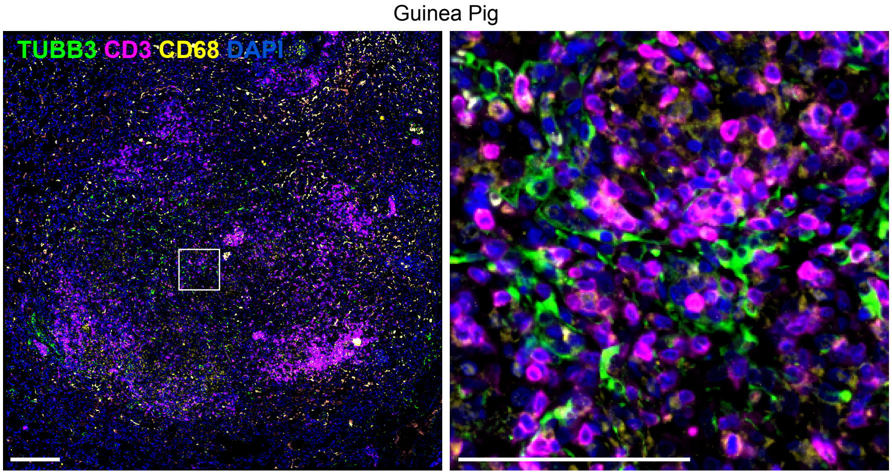

### Supplemental Figure 8

##### Monard, et al. Supplemental Figure 8

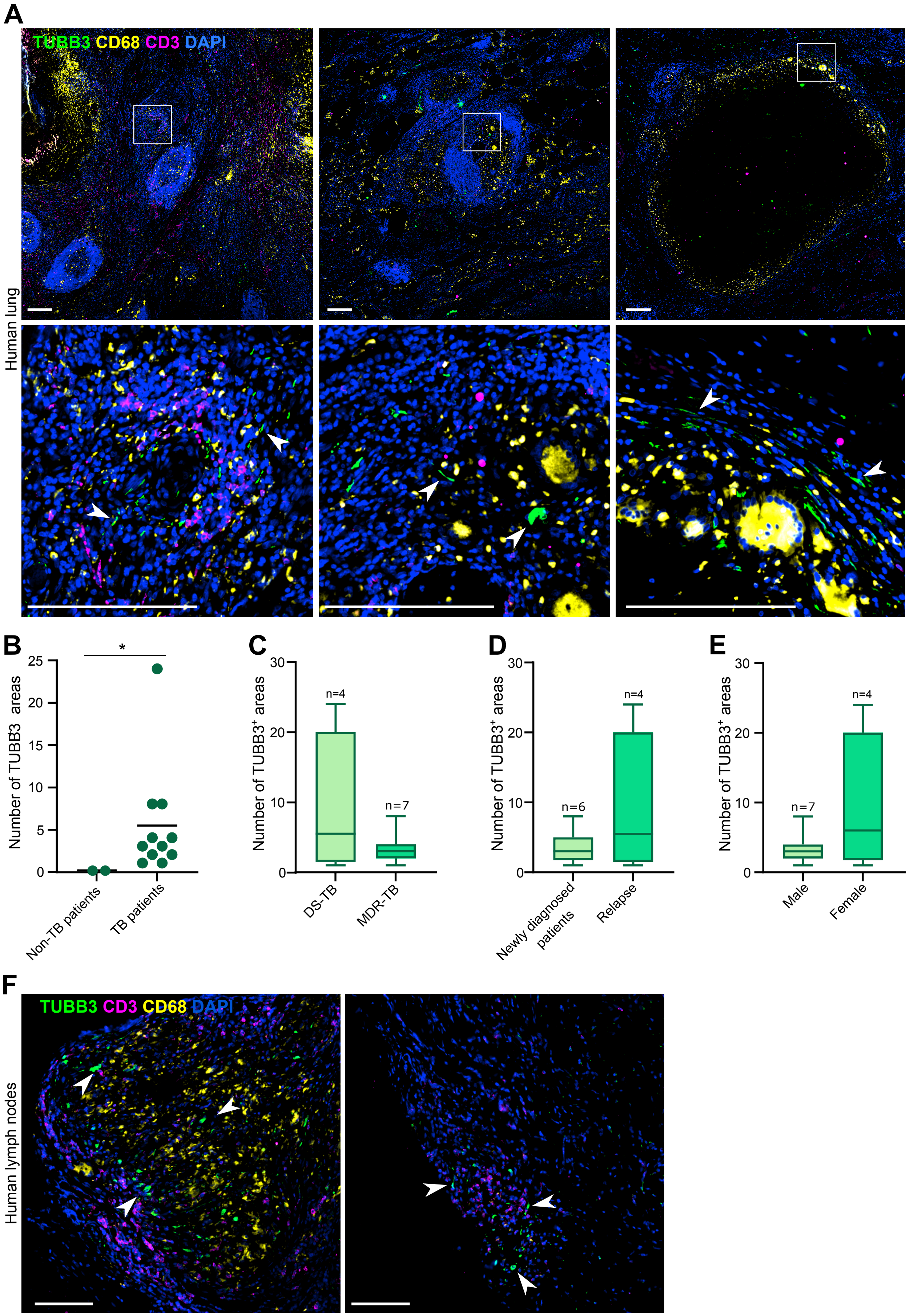
